# Reconstruction of the carbohydrate 6-O sulfotransferase gene family evolution in vertebrates reveals novel member, CHST16, lost in amniotes

**DOI:** 10.1101/667535

**Authors:** Daniel Ocampo Daza, Tatjana Haitina

## Abstract

Glycosaminoglycans are sulfated polysaccharide molecules, essential for many biological processes. The 6-O sulfation of glycosaminoglycans is carried out by carbohydrate 6-O sulfotransferases (C6OST), previously named Gal/GalNAc/GlcNAc 6-O sulfotransferases. Here for the first time we present a detailed phylogenetic reconstruction, analysis of gene synteny conservation and propose an evolutionary scenario for the C6OST family in major vertebrate groups, including mammals, birds, non-avian reptiles, amphibians, lobe-finned fishes, ray-finned fishes, cartilaginous fishes and jawless vertebrates. The C6OST gene expansion likely occurred in the chordate lineage, after the divergence of tunicates and before the emergence of extant vertebrate lineages.

The two rounds of whole genome duplication in early vertebrate evolution (1R/2R) only contributed two additional C6OST subtype genes, increasing the vertebrate repertoire from four genes to six, divided into two branches. The first branch includes *CHST1* and *CHST3* as well as a previously unrecognized subtype, *CHST16*, that was lost in amniotes. The second branch includes *CHST2*, *CHST7* and *CHST5*. Subsequently, local duplications of *CHST5* gave rise to *CHST4* in the ancestor of tetrapods, and to *CHST6* in the ancestor of primates.

The teleost specific gene duplicates were identified for *CHST1*, *CHST2* and *CHST3* and are result of whole genome duplication (3R) in the teleost lineage. We could also detect multiple, more recent lineage-specific duplicates. Thus, the repertoire of C6OST genes in vertebrate species has been shaped by different events at several stages during vertebrate evolution, with implications for the evolution of the skeleton, nervous system and cell-cell interactions.

## Introduction

Glycosaminoglycans (GAG) are sulfated linear polysaccharide molecules comprised of repeating disaccharides, like chondroitin sulfate (CS), dermatan sulfate (DS), keratan sulfate (KS) and heparan sulfate (HS). CS and DS are composed of N-acetylgalactosamine (GalNAc) linked to glucuronic acid or iduronic acid, respectively. HS and KS are composed of N-acetylglucosamine (GlcNAc) linked to glucuronic acid or galactose, respectively. Sulfated GAGs are found in both vertebrates and invertebrates and are important for many biological processes like cell adhesion, signal transduction and immune response (Yamada et al. 2011; Soares da Costa et al. 2017). The polymerization of long, linear GAG chains onto core protein takes place in the Golgi apparatus and results in the formation of proteoglycans that are essential components of the extracellular matrix (Kjellén & Lindahl 1991).

Sulfation is a complex posttranslational modification process that is common for GAGs and is important for their activity (Soares da Costa et al., 2017). The 6-O sulfation of CS and KS is carried out by enzymes of the carbohydrate 6-O sulfotransferase (C6OST) family (Kusche-Gullberg & Kjellén 2003). The nomenclature of the reported members of this family is summarized in table 1. Hereafter we will use the abbreviation C6OST to refer to the protein family in general and the symbols *CHST1* through *CHST7* to refer to the individual genes. The 6-O sulfation of HS is carried out by enzymes of another protein family, the heparan sulfate 6-O sulfotransferases (HS6ST1-HS6ST3) (Nagai & Kimata 2014) that will not be discussed here (Kusche-Gullberg & Kjellén 2003). The sulfation of GAGs is carried out by using 3’-phosphoadenosine 5’-phosphosulfate (PAPS) as a sulfonate donor. PAPS binds to C6OSTs at specific sequence motifs: the 5’-phosphosulfate binding (5’PSB) motif RxGSSF (Habuchi et al. 2003) and the 3’-phosphate binding (3’PB) motif RDPRxxxxSR (Tetsukawa et al. 2010). As an example, in the human chondroitin 6-sulfotransferase 1 protein sequence (encoded by *CHST3*), the 5’PSB motif corresponds to positions 142-147 (RTGSSF), and the 3’PB motif to positions 301-310 (RDPRAVLASR). Chondroitin 6-O sulfotransferase 1 is one of the most widely studied members of the C6OST family, showing activity towards both CS and KS (see references in Yusa et al. 2006). Another enzyme, chondroitin 6-O sulfotransferase 2 (encoded by *CHST7*) also shows activity towards CS (Kitagawa et al. 2000). The expression of *CHST3* and *CHST7* has been reported in a number of tissues, including chondrocytes, immune system organs, as well as the central and peripheral nervous system (see Habuchi (2014) and references therein). Mutations in the *CHST3* gene are associated with congenital spondyloepiphyseal dysplasia, congenital joint dislocations and hearing loss in kindred families (Srivastava et al. 2017; Waryah et al. 2016), whereas overexpression of *CHST3* lowers the abundance of the proteoglycan aggrecan in the extracellular matrix of the aged brain, which is associated with the loss of neural plasticity (Miyata & Kitagawa 2016). The orthologues of *CHST3* and *CHST7* have been characterized in zebrafish, where *chst3a*, *chst3b* and *chst7* are expressed in pharyngeal cartilages, the notochord and several brain regions during development (Habicher et al. 2015).

**Table 1.**
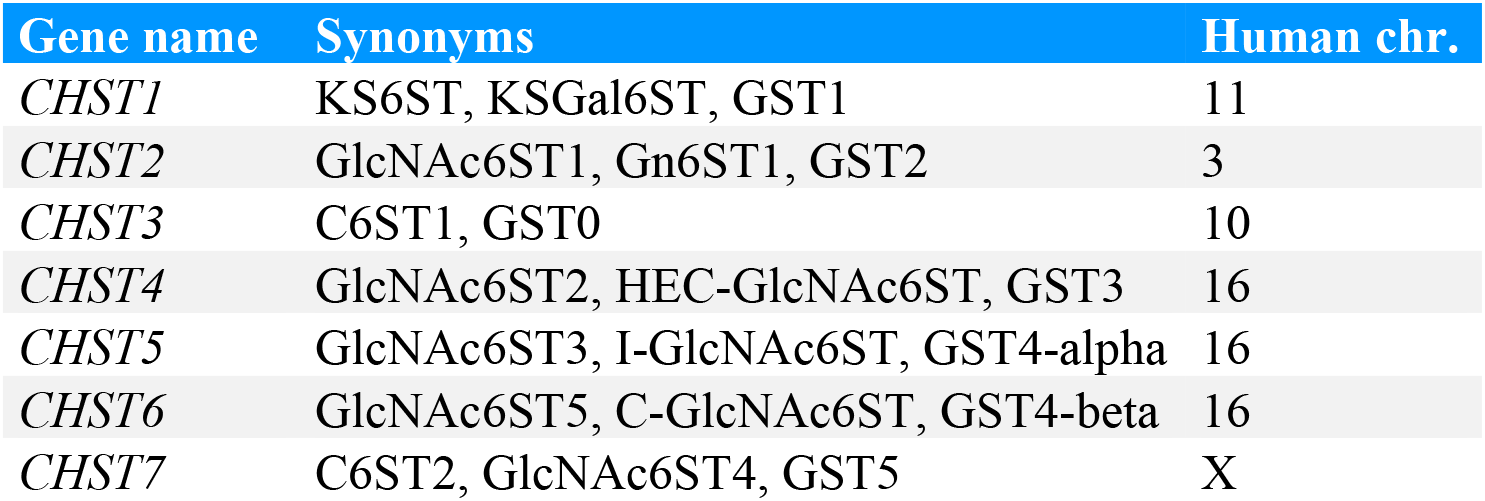
Nomenclature of the carbohydrate 6-O sulfotransferases.

Carbohydrate Sulfotransferase 1 (encoded by *CHST1*), also called Keratan Sulfate Gal-6 Sulfotransferase 1 (KSGal6ST), is responsible for the sulfation of KS. *CHST1* expression has been described in the developing mouse brain (Hoshino et al. 2014) and in human endothelial cells (Li & Tedder 1999). Another sulfotransferase showing activity towards keratan sulphate is the corneal N-acetylglucosamine-6-O-sulfotransferase (C-GlcNAc6ST, encoded by *CHST6*). In humans, the closely related *CHST5* and *CHST6* genes are located approximately 40 Kb apart on chromosome 16. Whereas in the mouse genome there is only one gene in the corresponding chromosomal region on chromosome 8, called *CHST5*. While human *CHST5* expression seems restricted to the small intestine and colon (Lee et al. 1999), mutations of *CHST6* in humans cause a rare autosomal recessive macular corneal dystrophy (Akama et al. 2000; Carstens et al. 2016; Rubinstein et al. 2016). In mice, a similar condition in the form of CS/DS aggregates in the cornea is caused by the disruption of the *CHST5* gene (Parfitt et al. 2011). In addition, the similarity in enzymatic activity between the human *CHST6* and mouse *CHST5* gene products has been used to suggest that these genes are orthologous (Akama et al. 2001, 2002).

N-acetylglucosamine-6-O-sulfotransferase 1 (GlcNAc6ST1, encoded by *CHST2*) is expressed in brain tissues and several internal organs of adult mice (Fukuta et al. 1998). It has also been reported in endothelial tissues of both humans and mice (Li & Tedder 1999). In the mouse model, *CHST2* is critical for neuronal plasticity in the developing visual cortex (Takeda-Uchimura et al. 2015), and *CHST2* deficiency leads to increased levels of amyloid-β phagocytosis thus modulating Alzheimer’s pathology (Zhang et al. 2017). N-acetylglucosamine-6-O-sulfotransferase 2 (GlcNAc6ST2, encoded by *CHST4*) is expressed exclusively in the high endothelial venules of the lymph nodes (Bistrup et al. 2004) and its disruption in mice affects lymphocyte trafficking (Hemmerich, Bistrup et al. 2001). *CHST4* is also expressed in early-stage uterine cervical and corpus cancers (Seko et al. 2009).

Functional studies of GAGs have expanded greatly during the last twenty years. The roles of GAG modifying enzymes, including C6OSTs, in health and disease have been studied extensively in human and mouse, as well as to some extent in chicken (Fukuta et al. 1995; Yamamoto et al. 2001; Nogami et al. 2004; Kobayashi et al. 2010). However, there is a big information gap regarding which C6OST genes can be found outside mammalian vertebrates, as well as the phylogenetic relationship between them. Outside mammals and chicken, the number of studies is very limited to just a few species, including zebrafish (*Danio rerio*) (Habicher et al. 2015), as well as vase tunicate (*Ciona intestinalis*) (Tetsukawa et al. 2010), and pearl oyster (*Pinctada fucata martensii*) (Du et al. 2017).

Here for the first time we describe the evolution of the C6OST family of sulfotransferases based on detailed phylogenetic and chromosomal location analyses in major vertebrate groups, and propose a duplication scenario for the expansion of the C6OST family during vertebrate evolution. We also report a previously unrecognized KSGal6ST-like subtype of C6OST, encoded by a gene we have called *CHST16*, that is lost in amniotes.

## Materials and Methods

### Identification of C6OST gene sequences

C6OST amino acid sequences corresponding to the *CHST1-7* genes were sought primarily in genome assemblies hosted by the National Center for Biotechnology Information (NCBI) Assembly database (www.ncbi.nlm.nih.gov/assembly) (Kitts et al. 2016; Canese et al. 2017) and the Vertebrate Genomes Project (VGP, vertebrategenomesproject.org) (The G10K consortium, *manuscript in preparation*). For some species, the Ensembl genome browser (www.ensembl.org) (Perry et al. 2018) was used. C6OST sequences were also sought in transcriptome assemblies hosted by the PhyloFish Portal (phylofish.sigenae.org) (Pasquier et al. 2016). Independent sources for an Eastern Newt reference transcriptome (Abdullayev et al. 2013) and the gulf pipefish genome assembly (Small et al. 2016) were also used. All the investigated species, with their binomial nomenclature, and genome/transcriptome assemblies are listed in supplementary table S1 together with links to their sources. In total, 158 species were investigated, representing the major vertebrate groups. They include 18 mammalian species; 33 avian species in 16 orders; 14 non-avian reptile species; six amphibian species; the basal lobe-finned fish coelacanth; the holostean fishes spotted gar and bowfin; 67 teleost fish species in 35 orders, including the important model species zebrafish; seven cartilaginous fish species; as well as three jawless vertebrate species, the sea lamprey, Arctic lamprey (also known as Japanese lamprey), and inshore hagfish. C6OST sequences from invertebrate species were also sought. Notably, the vase tunicate was used to provide a relative dating point with respect to the early vertebrate whole genome duplications (1R/2R), and the fruit fly was used as an outgroup.

The C6OST sequences were identified by searching for NCBI Entrez Gene models (Maglott et al. 2011; Brown et al. 2014) or Ensembl gene predictions (Curwen et al. 2004) annotated as *CHST1*-7, followed by extensive *tblastn* searches (Altschul et al. 1990) to identify sequences with no corresponding gene annotations or for species where gene annotations were not available. For invertebrate species, sequences were also sought using the profile-Hidden Markov Model search tool HMMER (hmmer.org) (Finn et al. 2011), aimed at reference proteomes. In most cases (excluding transcriptome data), corresponding gene models could be found and their database IDs were recorded. In cases where no gene models could be identified, or where they included errors (Prosdocimi et al. 2012), C6OST sequences were predicted/corrected by manual inspection of their corresponding genomic regions, including flanking regions and introns. Exons were curated with respect to consensus sequences for splice donor/acceptor sites (Rogozin et al. 2012; Abril et al. 2005) and start of translation (Nakagawa et al. 2008), as well as sequence homology to other C6OST family members.

All identified amino acid sequences were collected and the corresponding genomic locations (if available) were recorded. All collected sequences were verified against the Pfam database of protein families (pfam.xfam.org) (El-Gebali et al. 2019) to ensure they contained a sulfotransferase type 1 domain (Pfam ID: PF00685), and inspected manually to verify the characteristic 5’-phosphosulfate binding (5’PSB) and 3’-phosphate binding (3’PB) motifs. All genomic locations and sequence identifiers used in this study have been verified against the latest genome assembly and database versions, including NCBI’s Reference Sequence Database (RefSeq 92, 8 January 2019) and Ensembl (version 95, January 2019).

### Sequence alignment and phylogenetic analyses

Sequence alignments were constructed with the MUSCLE alignment algorithm (Edgar 2004) applied through AliView 1.25 (Larsson 2014). Alignments were curated manually in order to adjust poorly aligned stretches with respect to conserved motifs and exon boundaries, as well as to identify faulty or incomplete sequences. Phylogenies were constructed using IQ-TREE v1.6.3 which applies a stochastic maximum likelihood algorithm (Nguyen et al. 2015). The best-fit amino acid substitution model and substitution parameters were selected using IQ-TREE’s model finder with the -m TEST option (Kalyaanamoorthy et al. 2017). The proportion of invariant sites was optimized using the --opt-gamma-inv option. Branch supports were calculated using IQ-TREE’s non-parametric ultrafast bootstrap (UFBoot) method (Minh et al. 2013) with 1000 replicates, as well as the approximate likelihood-ratio test (aLRT) with SH-like supports (Anisimova & Gascuel 2006; Anisimova et al. 2011) over 1000 iterations.

### Conserved synteny analyses

The two whole genome duplications that occurred at the base of vertebrate evolution (1R and 2R) resulted in a large number of quartets of related chromosome regions, each such quartet is called a paralogon or a paralogy group, and related chromosome regions are said to be paralogous. To investigate whether any of the vertebrate C6OST genes arose in the 1R/2R whole genome duplication events, and duplicated further in the teleost whole genome duplication 3R, we attempted to detect known paralogous chromosome regions around C6OST genes. This was done through the detection of conserved synteny, the conservation of gene family co-localizations across different chromosome regions that reflects the duplications of not only individual genes but larger chromosome segments and by extension the whole genome. We carried out a conserved synteny analysis around the C6OST gene-bearing chromosome regions in the human, Carolina anole lizard, spotted gar and zebrafish genomes. The anole lizard was chosen because of the presence of a “*CHST4*/*5*-like” gene in this species. The spotted gar was chosen because of the presence of *CHST16*, which is missing from amniotes, because it shows a moderate degree of genome rearrangement compared with teleost fish genomes, and because its taxonomic position allows it to bridge the gap between ray-finned fishes and lobe-finned fishes (including tetrapods) (Amores et al. 2011; Braasch et al. 2016). The zebrafish genome was chosen to investigate the involvement of 3R. Lists of gene predictions 2.5 Mb upstream and downstream of the C6OST genes in these species were downloaded using the BioMart function in Ensembl version 83. These lists were sorted according to Ensembl protein family predictions (Ensembl 83 is the last version of the database to use these family predictions) in order to identify gene families with members on at least two of the C6OST gene-bearing chromosome blocks within each species. Sequence identifiers and genomic locations for each member gene in these gene families were collected for the following species: human, chicken, Western clawed frog, spotted gar, zebrafish, medaka, and elephant shark to identify regions that show ancestral paralogy relationships. All locations and sequence identifiers were verified against the latest versions of the genome assemblies in the NCBI Assembly database.

Only one *CHST1* gene could be identified in the zebrafish (see results below). Therefore, to investigate the conserved synteny between duplicated *CHST1* genes in teleost fishes, the regions of the channel catfish *CHST1* genes on chromosomes 4 and 8 were also analysed. Gene models 1 Mb to each side of the channel catfish *CHST1* genes were identified in the NCBI Genome Data Viewer and used to identify the homologous chromosome regions in the zebrafish genome. These regions were then used to identify gene families with members on both chromosome regions through Ensembl’s BioMart, the same way as described above.

## Results

### Phylogeny of the C6OST family

C6OST amino acid sequences were collected from a large diversity of vertebrate genomes and aligned in order to produce phylogenies of the C6OST gene family across vertebrates. The first phylogeny of C6OST family in chordates is summarized in figure 1 and includes the vase tunicate along with a smaller subset of representative vertebrates. This phylogeny is rooted with two putative family members from the fruit fly, *CG31637* and *CG9550*. These sequences share the sulfotransferase domain (Pfam ID: PF00685) and recognizable 5’PSB and 3’PB motifs with the vertebrate C6OST sequences. In addition to this phylogeny, we produced detailed phylogenies for each C6OST subtype branch including all investigated vertebrate species. These have been included as supplementary figures S1-S6. This was done in order to classify all identified sequences into correct C6OST subtype, and to improve the resolution of some phylogenetic relationships that were unclear in the smaller phylogeny.

**Fig. 1.**
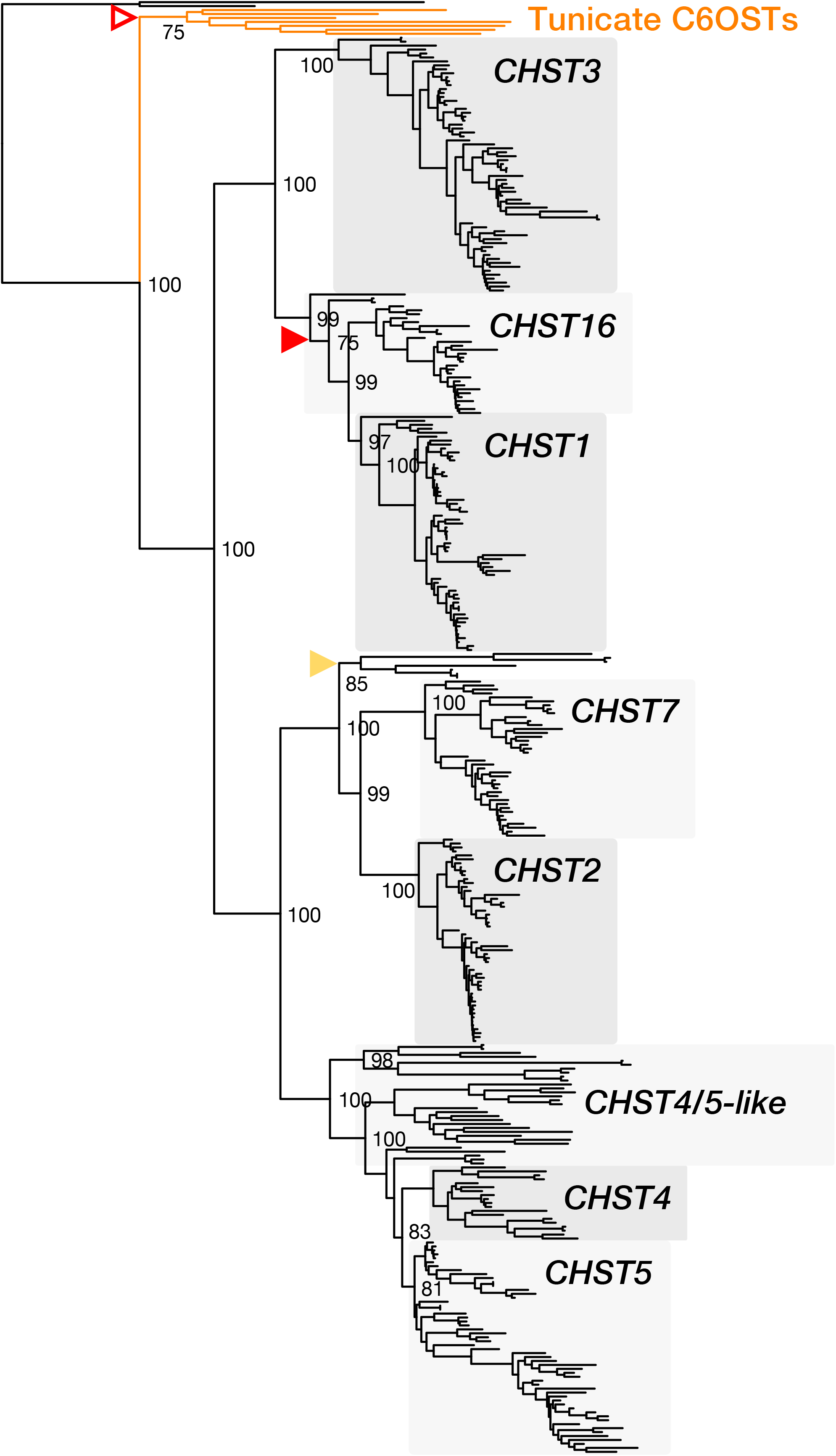
Maximum likelihood phylogeny of chordate C6OST sequences. The phylogeny is supported by approximate Likelihood-Ratio Test (aLRT) and UltraFast Bootstrap (UFBoot) analyses. UFBoot supports for deep nodes are shown. Red filled arrowheads indicate unsupported nodes (≤ 75%) in both UFBoot and aLRT, yellow filled arrowheads indicate nodes with low aLRT support only, red open arrowheads indicate nodes with low UFBoot support only. The phylogeny is rooted with the *Drosophila melanogaster* C6OST sequences *CG9550* and *CG31637*.

Our phylogeny (figure 1) supports the subdivision of the C6OST family into two main branches. The first branch contains three well-supported subtype clades, including *CHST1* and *CHST3* as well as a previously unrecognized *CHST1*-like subtype of genes we have named *CHST16*. The second C6OST branch in vertebrates contains well-supported *CHST2*, *CHST4*, *CHST5* and *CHST7* clades, as well as several smaller clades of *CHST4*/*5-like* sequences from jawless vertebrates (*Agnatha*), amphibians and non-avian reptiles. We could identify *CHST16*, *CHST3*, as well as *CHST2*/*7-like* and *CHST4*/*5-like* sequences in lampreys (described in detail below), as well as *CHST1*, *CHST16*, *CHST2*/*7-like* and *CHST4*/*5-like* sequences in the inshore hagfish. While the identity of some of these sequences remains unclear, the jawless vertebrate branches place the divergences between the C6OST subtype clades before the split between jawless and jawed vertebrates (*Gnathostomata*), early in vertebrate evolution. To provide an earlier dating point, we used seven C6OST-like sequences from the vase tunicate. These sequences were first identified by Tetsukawa et al. (2010), and our phylogeny (figure 1) supports their conclusion that the tunicate C6OST genes represent an independent lineage-specific gene expansion. Thus, the diversification of vertebrate C6OST genes likely occurred in a chordate or vertebrate ancestor after the divergence of tunicates, that took place 518-581 Mya (Hedges and Kumar, 2009), This is consistent with the time window of the two rounds of whole-genome duplication early in vertebrate evolution, 1R/2R (Dehal & Boore 2005; Nakatani et al. 2007; Putnam et al. 2008; Sacerdot et al. 2018). We could also identify teleost-specific duplicates of *CHST1*, *CHST2*, and *CHST3* (described below), which is consistent with the third round of whole-genome duplication (3R) that occurred early in the teleost lineage (Meyer & van de Peer 2005).

### *CHST1* and *CHST16*

The *CHST1* and *CHST16* branches of the C6OST phylogeny are shown in figure 2A. The full-species phylogeny of *CHST1* sequences is shown in supplementary figure S1, and of *CHST16* sequences in supplementary figure S2. Both *CHST1* and *CHST16* are represented in cartilaginous fishes (*Chondrichthyes*), lobe-finned fishes (*Sarcopterygii*), including tetrapods and coelacanth, as well as ray-finned fishes (*Actinopterygii*), including the spotted gar and teleost fishes. Overall, both *CHST1* and *CHST16* branches follow the accepted phylogeny of vertebrate groups, and the overall topology is well-supported. However, there are some notable inconsistencies: We could identify putative *CHST1* and *CHST16* orthologs in the inshore hagfish, as well as putative *CHST16* orthologs in the Arctic and sea lampreys. However, the jawless vertebrate *CHST16* sequences cluster basal to both jawed vertebrate *CHST1* and *CHST16* clades. Nevertheless, our overall results indicate that both *CHST1* and *CHST16* were present in a vertebrate ancestor prior to the split between jawless and jawed vertebrates. Within teleost fishes, we could identify duplicate *CHST1* genes located on separate chromosomes, which we have named *CHST1a* and *CHST1b*. However, the spotted gar *CHST1* sequence clusters together with the teleost *CHST1a* clade rather than basal to both *CHST1a* and *CHST1b* clades. This is true for the smaller phylogeny shown in figure 2A as well as for the phylogeny with full species representation (supplementary figure S1), which also includes a *CHST1* sequences from another holostean fish, the bowfin. The duplicate *CHST1* genes in the investigated eel species also *both* cluster within the *CHST1a* branch. Thus, we could not determine with any certainty whether these *CHST1* duplicates in eels represent *CHST1a* and *CHST1b*. These inconsistencies are likely, at least partially, caused by uneven evolutionary rates between different lineages as well as a relatively high degree of sequence conservation for *CHST1* sequences. There are also duplicate *CHST1*, but not *CHST16*, genes in the allotetraploid African clawed frog, whose locations on chromosomes 4L and 4S correspond to each of the two homeologous sub-genomes (Session et al. 2016).

**Fig. 2.**
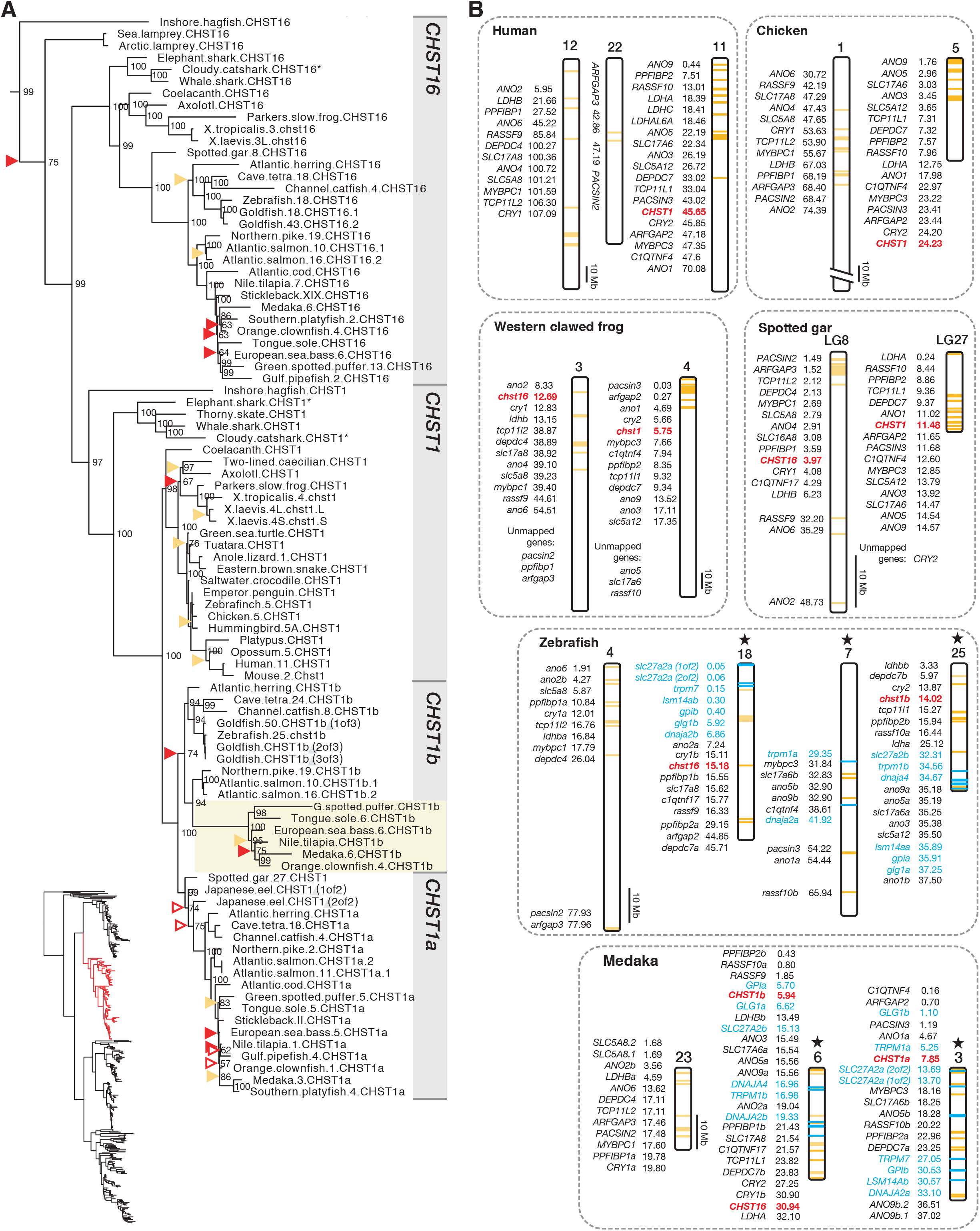
**A.** Phylogeny of the *CHST1* and *CHST16* branch of C6OST sequences. The placement of the branch within the full C6OST phylogeny is indicated in the bottom left. Sequence names include species names followed by chromosome/linkage group designations (if available) and gene symbols. Asterisks indicate incomplete sequences. For node support details, see figure 1 caption. Some node support values for shallow nodes have been omitted for visual clarity. The fast-evolving branch of neoteleost *CHST1b* sequences is highlighted. **B.** Conserved synteny between *CHST1* and *CHST16*-bearing chromosome regions in tetrapods and the spotted gar, including *CHST1a*, *CHST1b*, and *CHST16*-bearing chromosome regions in teleost fishes. Stars indicate chromosome regions in common between two different conserved synteny analyses: *CHST1* and *CHST16*-bearing regions (yellow labels), and *CHST1a* and *CHST1b*-bearing regions (blue labels).

There have been notable gene losses of both *CHST1* and *CHST16*. *CHST16* genes could not be identified in non-avian reptiles, birds or mammals, indicating that this subtype was lost in the amniote ancestor. Within cartilaginous fishes, *CHST16* was missing in all investigated skate species, indicating a loss from *Rajiformes* and possibly more generally from *Batoidea*. *CHST16* genes were also missing from both eel species, indicating a loss within the genus *Anguilla*, and possibly more generally within *Elopomorpha*. Not all teleost fish species preserve both *CHST1* duplicates. Notably, *CHST1a* is missing from cypriniform fishes, including the zebrafish, and we could not identify *CHST1a* in one salmoniform fish, the Arctic char. The *CHST1b* gene seems to have been lost more widely: We could not find *CHST1b* sequences in several basal spiny-rayed fishes (*Acanthomorpha*), such as opah (order *Lampriformes*), Atlantic cod (order *Gadiformes*), and longspine squirrelfish (order *Holocentriformes*), as well as several basal percomorph fishes, namely bearded brotula (order *Ophidiiformes*), mudskippers (order *Gobiiformes*) which instead have duplicate *CHST1a* genes, yellowfin tuna (order *Scombriformes*), tiger tail seahorse and gulf pipefish (order *Syngnathiformes*). It was also missing from turbot (order *Pleuronectiformes*), turquoise killifish, guppy and Southern platyfish (order *Cyprinodontiformes*), barred knifejaw (order *Centrarchiformes*), European perch and Three-spined stickleback (order *Perciformes*), and Japanese pufferfish (order *Tetraodontiformes*). At least some of these losses could be due to incomplete genome assemblies, as there are closely related species that do have both *CHST1a* and *CHST1b* genes, such as the giant oarfish (order *Lampriformes*), blackbar soldierfish (order *Holocentriformes*), tongue sole (order *Pleuronectiformes*), Murray cod (order *Centrarchiformes*), tiger rockfish (order *Perciformes*), and green spotted pufferfish (order *Tetraodontiformes*). We have followed the phylogeny and classification of teleost fishes by Near et al. (2012) and Betancur-R et al. (2017). Aside from this diversity in terms of *CHST1b* gene preservation or loss, *CHST1b* genes seem to have evolved more rapidly in neoteleosts, as indicated by the branch lengths within this clade in our phylogenies (highlighted in figure 2A and supplementary figure S1). The neoteleost *CHST1b* genes also have a divergent exon structure (see *“Exon/intron structures of C6OST genes”* below).

We could identify 15 gene families with members in the vicinity of both *CHST1* and *CHST16* (subset 1, supplementary table S2): ANO1/2, ANO3/4/9, ANO5/6/7, ARFGAP, C1QTNF4/17, CRY, DEPDC4/7, LDH, MYBPC, PACSIN, PPFIBP, RASSF9/10, SLC5A5/6/8/12, SLC17A6/8, and TCP11. This was detected in the spotted gar genome on linkage groups 27 and 8 (figure 2B). All identified chromosome segments are shown in supplementary data 3. In the human genome, the identified blocks of conserved synteny correspond mainly to segments of chromosomes 11 (where *CHST1* is located), 12 and 22 (figure 2B), as well as 1, 6, 19 and X (not shown here). These chromosome regions have been recognized as paralogous, the result of the 1R/2R whole genome duplications, in several large-scale reconstructions of vertebrate ancestral genomes (Nakatani et al. 2007; Putnam et al. 2008; Sacerdot et al. 2018). The *CHST1* and *CHST16*-bearing chromosome regions correspond to the “D” paralogon in Nakatani et al. (2007), more specifically the vertebrate ancestral paralogous segments called “D1” (*CHST1*) and “D0” (*CHST16*). In the study by Sacerdot et al. (2018), they correspond to the reconstructed pre-1R ancestral chromosome 6. One of the authors has previously analyzed these chromosome regions extensively, and could also conclude that they arose in 1R/2R (Lagman et al. 2013; Ocampo Daza & Larhammar 2018).

With respect to 3R, we could identify seven neighboring gene families in subset 1 which also had teleost-specific duplicate genes in the vicinity of *CHST1a* and *CHST1b*: ANO1/2, ANO3/4/9, ANO5/6/7, DEPDC4/7, SLC17A6/8, PPIFBP, and RASSF9/10 (figure 2B). We could also identify a further six gene families with members in the vicinity of *CHST1a* and *CHST1b* in teleosts (subset 2, supplementary table S2): DNAJA1/2/4, GLG1, GPI, LSM14A, SLC27A, and TRPM1/3/6/7 (labeled with blue color on figure 2B). The identified conserved synteny blocks correspond to segments of chromosomes 18 and 7 on one side, and 25 on the other, in the zebrafish genome, as well as segments of chromosomes 6 and 3 in the medaka genome. These chromosome segments have previously been identified to be the result of the teleost specific whole-genome duplication, 3R (Kasahara et al. 2007; Nakatani et al. 2007; Nakatani & McLysaght 2017). In the reconstruction of the pre-3R genome by Nakatani & McLysaght (2017), these regions correspond to proto-chromosome 10. The chromosome segments we identified in the spotted gar, chicken and human genome also agree with the reconstruction by Nakatani & McLysaght (2017). In the analyses of teleost fish chromosome evolution by Kasahara et al. (2007), as well as in (Nakatani et al. 2007), these regions correspond to proto-chromosome “j”. We could also detect extensive translocations between paralogous chromosome segments in teleost fishes that originated in 1R/2R. See chromosome 18 in the zebrafish and chromosome 6 in the medaka, as an example (figure 2B). This is also compatible with previous observations (Ocampo Daza & Larhammar 2018; Kasahara et al. 2007; Nakatani & McLysaght 2017). These homology-related chromosome rearrangements likely underlie the co-localization of *CHST16* with *CHST1b* in many teleost genomes, as on chromosome 6 in the medaka (figure 2B), or with *CHST1a* within *Otocephala*, as on chromosome 18 in the Mexican cave tetra (see chromosome designations in the phylogeny in figure 2A). Cypriniform fishes, including the zebrafish, have lost the *CHST1a* gene as described above. However, the *CHST16* gene is located near the chromosome segment where *CHST1a* would have been located in the zebrafish genome (figure 2B). The co-localization of *CHST16* with either *CHST1a* or *CHST1b* occurs only within teleost fishes, and could be detected in all investigated teleost genomes that have been mapped to chromosomes or assembled into linkage groups (supplementary data S1).

In addition to 3R, there have been more recent whole-genome duplication events within cyprinid fishes (family *Cyprinidae*) and salmoniform fishes (order *Salmoniformes*) (Glasauer & Neuhauss 2014). We identified additional duplicates of *CHST1b* in the goldfish genome, however only one of the three duplicates is mapped, to chromosome 50 (figure 2A), and the other two are identical. There are also duplicates of the *CHST16* gene on goldfish chromosomes 18 and 43. These chromosome locations correspond to homeologous chromosomes that arose in the cyprinid-specific whole-genome duplication (see figure 2 in Chen et al. 2018). In salmoniform fishes, there are additional duplicates of *CHST1a* in the Atlantic salmon and of *CHST1b* and *CHST16* in all three investigated species, Atlantic salmon, rainbow trout and Arctic char. While several of these salmoniform duplicates are unmapped, including one of the *CHST1a* duplicates in Atlantic salmon, the chromosomal locations of the syntenic *CHST1b* and *CHST16* duplicates on chromosomes 10 and 16 in Atlantic salmon (figure 2 in Lien et al.2016), 2 and 1 in rainbow trout (figure 2 in Berthelot et al. 2014), as well as 4 and 26 in Arctic char (figure 1 in Christensen et al. 2018), correspond to duplicated chromosome segments from the salmoniform-specific whole-genome duplication.

### CHST3

The *CHST3* branch of our C6OST phylogeny is shown in figure 3A. The full-species phylogeny of *CHST3* sequences is shown in supplementary figure S3. We could identify *CHST3* sequences across all major vertebrate lineages, including both lamprey species but not the inshore hagfish. The allotetraploid African clawed frog has duplicated *CHST3* genes located on chromosomes 7L and 7S that correspond to each of the two homeologous sub-genomes (Session et al. 2016). Overall, our phylogenies follow the accepted phylogeny of vertebrate groups. However, following are some inconsistencies in both phylogenies. The cartilaginous fish clade is not resolved, however all cartilaginous fish *CHST3* sequences cluster basal to the bony vertebrate clade. In addition, the coelacanth *CHST3* sequence clusters basal to both the tetrapod and ray-finned fish clades. Nonetheless, the overall topologies of our phylogenetic analyses support the presence of a *CHST3* gene before the split between jawless and jawed vertebrates.

**Fig. 3.**
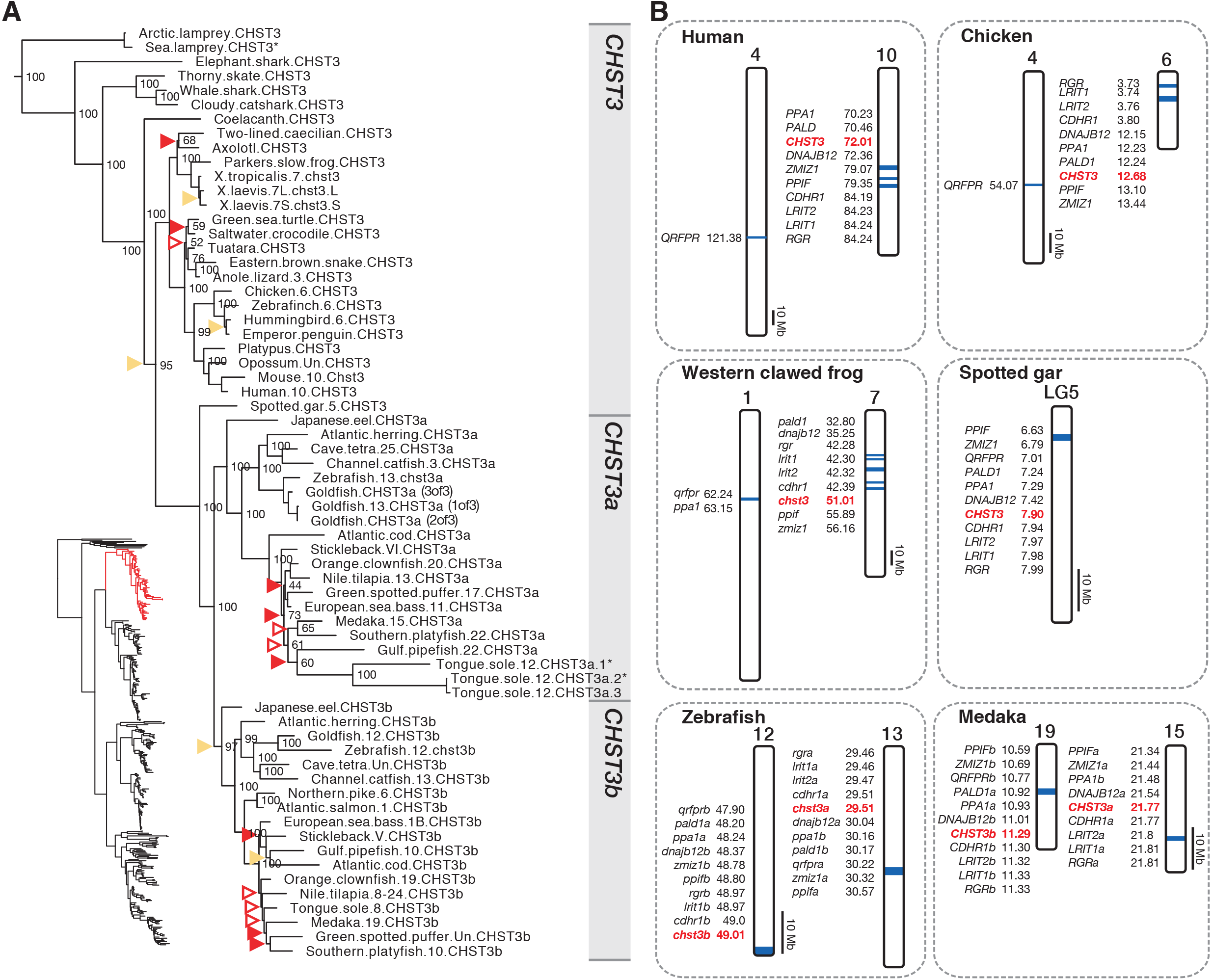
**A.** Phylogeny of the *CHST3* branch of C6OST sequences. See figure 2 caption for details. **B.** Conserved synteny across *CHST3*, *CHST3a* and *CHST3b*-bearing chromosome regions.

In teleost fishes we found duplicates of *CHST3* located on different chromosomes (figure 3B). These genes have been named *chst3a* and *chst3b* in the zebrafish (Habicher et al. 2015). The single *CHST3* sequence from spotted gar clusters at the base of the well-supported teleost *chst3a* and *chst3b* clades, which is consistent with duplication in the time window of 3R. We could identify nine gene families with teleost-specific duplicate members in the vicinity of *chst3a* and *chst3b*: CDHR1, DNAJB12, LRIT, PALD1, PPA, PPIF, QRFPR, RGR, and ZMIZ (subset 3, supplementary table S2). The identified paralogous segments on chromosomes 13 and 12 in zebrafish and on chromosomes 15 and 19 in medaka (figure 3B), correspond to chromosome segments that most likely arose in 3R (Kasahara et al. 2007; Nakatani et al. 2007; Nakatani & McLysaght 2017). In Kasahara et al. (2007) and Nakatani et al. (2007) they correspond to pre-3R proto-chromosome “d”, and in Nakatani & McLysaght (2017) they correspond to proto-chromosome “4”. The chromosome segments we could identify in the spotted gar, chicken and human genomes (figure 3B) are also compatible with these previous studies, as well as with comparative genomic analyses of the spotted gar genome vs. the human and zebrafish genomes (see figures 3 and 4, Table S2 in Amores et al. 2011). Other smaller synteny blocks not shown here are shown in supplementary data 3. We could identify three copies of *CHST3a* in the goldfish genome, however only one of them is mapped, to chromosome 13. Thus, it is not possible to attribute at least one of the duplications to the cyprinid-specific whole-genome duplication. There are also three copies of the *CHST3a* gene in the tongue sole genome, located in tandem on chromosome 12. In salmoniform fishes, no *CHST3a* gene could be found, indicating a loss of this gene, and no additional gene duplicates of *CHST3b* seem to have been preserved.

**Fig. 4.**
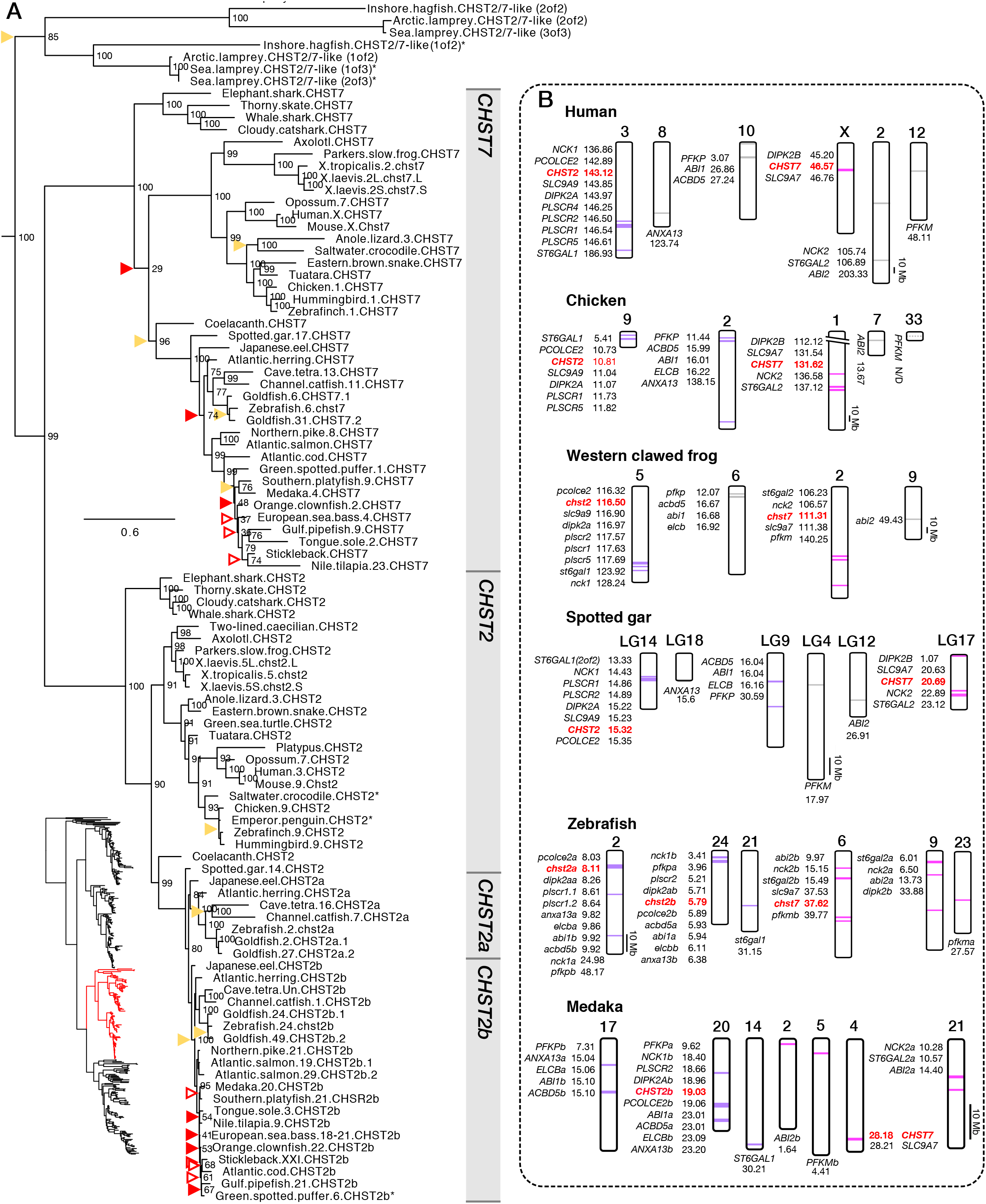
**A.** Phylogeny of the *CHST2* and *CHST7* branch of C6OST sequences. See figure 2 caption for details. **B.** Conserved synteny between *CHST2* and *CHST7*-bearing chromosome regions in tetrapods and spotted gar, including *CHST2a* and *CHST2b*-bearing regions in teleost fishes.

Our overall phylogeny of C6OST sequences shows that *CHST3* is closely related to *CHST1* and *CHST16* (figure 1). However, we could not identify any conserved synteny, i.e. no shared genomic neighbors, between *CHST3* and *CHST1* or *CHST16*, in any of the analyzed species. Based on our conserved synteny analysis, the *CHST1* and *CHST16* genes likely arose from an ancestral gene through 1R or 2R, as described above. This suggests not only that the *CHST3* gene was present together with the ancestral *CHST1/16* gene before 1R, but also that the *CHST3* and *CHST1/16* genes arose by a mechanism other than the duplication of a larger chromosome segment, as in a whole-genome duplication.

### *CHST2* and *CHST7*

The *CHST2* and *CHST7* branches of our C6OST phylogeny are shown in figure 4A. The full-species phylogeny of *CHST2* sequences is included in supplementary figure S4, and of *CHST7* sequences in supplementary figure S5. The *CHST2* and *CHST7* subtype branches are well-supported, and follow the accepted phylogeny of vertebrate groups, overall. Both clades include orthologs from lobe-finned fishes (including coelacanth and tetrapods), ray-finned fishes (including spotted gar and teleost fishes), as well as cartilaginous fishes. However, both coelacanth *CHST2* and *CHST7* cluster with the respective ray-finned fish clades rather than at the base of the lobe-finned fish clades. This is likely due to the low evolutionary rate, i.e. high degree of sequence conservation, of these coelacanth sequences (Amemiya et al. 2013) relative to the tetrapod sequences. There are duplicate *CHST2* and *CHST7* genes in the allotetraploid African clawed frog (*Xenopus laevis*), located on the homeologous chromosomes (Session et al. 2016) 5L and 5S, 2L and 2S, respectively.

Notably, *CHST7* could not be identified in any of the three investigated species within *Testudines* (turtles, tortoises and terrapins), indicating an early deletion of this gene within the lineage. *CHST7* is also missing from the two-lined caecilian (an amphibian) and several avian lineages, including penguins (order *Sphenisciformes*, three species were investigated), falcons (order *Falconiformes*, 2 species were investigated), as well as the rock pigeon (order *Columbiformes*), hoatzin (order *Ophistocomiformes*), downy woodpecker (order *Piciformes*), and hooded crow (order *Passeriformes*). For at least some of these species, the lack of a *CHST7* sequence could be due to incomplete genome assemblies, as there are other closely related species with *CHST7*, for example the band-tailed pigeon (order *Columbiformes*), and great tit (order *Passeriformes*) (supplementary data S3).

In jawless vertebrates, we could identify *CHST2* and *CHST7*-like sequences from the inshore hagfish, sea lamprey and Arctic lamprey genomes, forming two clades. These sequences cluster basal to both the jawed vertebrate *CHST2* and *CHST7* clades (figure 4A). Thus, we cannot determine whether these sequences represent either *CHST2* or *CHST7*, both *CHST2* and *CHST7*, or possibly an ancestral *CHST2*/*7* subtype.

There are duplicates of *CHST2* in teleost fishes located on different chromosomes (figure 4A). These have been named *chst2a* and *chst2b* in the zebrafish. We could identify both *chst2a* and *chst2b* only within basal teleost lineages (supplementary figure S4): eels (order *Anguilliformes*), freshwater butterflyfishes, bony-tongues and mormyrids (superorder *Osteoglossomorpha*), as well as *Otocephala*, including the Atlantic herring, European pilchard and allis shad (order *Clupeiformes*), zebrafish, Dracula fish, Amur ide and goldfish (order *Cypriniformes*), Mexican cave tetra and red-bellied piranha (order *Characiformes*), electric eel (order *Gymnotiformes*), and channel catfish (order *Siluriformes*). This indicates that the *CHST2a* gene was lost early in the euteleost lineage, which includes the majority of teleost species diversity. Both *CHST2a* and *CHST2b* clades are well-supported in the phylogeny (figure 4A), and the single spotted gar *CHST2* sequence clusters at the base of both branches, which is consistent with duplication within the time window of 3R. In the full-species phylogeny (supplementary figure S4) this topology is disrupted by the osteoglossomorph *CHST2a* and *CHST2b* branches. This is likely caused by uneven evolutionary rates for teleost *CHST2* sequences, as shown by the branch lengths within the phylogeny. Teleost *CHST2* sequences seem to have had a very low basal amino acid substitution rate overall, however the *CHST2a* sequences, as well as the *CHST2b* sequences in the African butterflyfish and mormyrids (superorder *Osteoglossomorpha*), seem to have evolved at a relatively faster rate. There are additional duplicates of both *CHST2a* and *CHST2b*, as well as of *CHST7*, in the goldfish genome, and their locations on chromosomes 2 and 27, 24 and 49, 6 and 31, respectively, supports their emergence through the cyprinid-specific whole genome duplication (see figure 2 in Chen et al. 2018). Salmoniform fishes lack *CHST2a*, as all euteleost fishes do, however there are duplicated *CHST2b* genes in the three species that were investigated. The *CHST2b* duplicates are located on chromosomes 19 and 29 in the Atlantic salmon genome, and 11 and 15 in the rainbow trout genome. These locations correspond to chromosomal segments known to have emerged in the salmoniform whole-genome duplication (Lien et al. 2016; Berthelot et al. 2014). In the Arctic char, only one of the *CHST2b* duplicates is mapped, to chromosome 14. No *CHST7* duplicates could be found in salmoniform fishes.

We could identify seven gene families with members in the vicinity of both *CHST2* and *CHST7*: BRINP, NOS1AP, PFK, and RGS4/5/8/16 were detected in the zebrafish genome, NCK and ST6GAL in the spotted gar genome, and SLC9A6/7/9 in both the spotted gar and human genomes (subset 4, supplementary table S2). Additionally, we could identify seven gene families with members in the vicinity of both *chst2a* and *chst2b* in the zebrafish genome: ABI, ACBD4/5, ANXA13, DIPK2, ELCB, PCOLCE, and PLSCR (subset 5, supplementary table S2). Out of both subsets, only four gene families, NCK, SLC9A6/7/9, ST6GAL, and DIPK2, had members in the vicinity of *CHST2* and *CHST7* in the tetrapod and/or spotted gar genomes. This apparent deficit of conserved syntenic genes possibly reflects a low gene density in the region of *CHST7*. We examined the chromosome regions around *CHST2* and *CHST7* genes in the synteny database Genomicus version 69.10 (http://www.genomicus.biologie.ens.fr/genomicus-69.10) (Muffato et al. 2010; Louis et al. 2015), and could only identify two gene pairs, in any species’ genome, in the vicinity of both *CHST2* and *CHST7*: *SLC9A9* and *SLC9A7*, as well as *C3orf58* and *CXorf36*. The latter gene pair was not identified in our synteny analysis. Nevertheless, the conserved synteny blocks that we could identify (figure 4B) correspond to chromosome regions recognized to have resulted from the 1R/2R whole genome duplications. In the reconstruction by Nakatani et al. (2007), these chromosome segments correspond to paralogon “F”, specifically the “F0” (*CHST2*) and “F3” (*CHST7*) vertebrate ancestral paralogous segments. In the analysis based around the Florida lancelet (*Branchiostoma floridae*) genome, they correspond to the reconstructed ancestral linkage group 10 (Putnam et al. 2008). In the more recent study by Sacerdot et al. (2018), they correspond to the reconstructed pre-1R ancestral chromosome 17. We could also identify one gene family, ITPR, with members in the vicinity of *chst7* and *chst16* in the zebrafish genome. However, this synteny pattern is not reproduced in any of the other investigated genomes (supplementary data S3).

With respect to 3R, seven gene families from subset 4 and subset 5 have teleost-specific duplicate members in the vicinity of both *chst2a* and *chst2b* in the zebrafish genome: PFK from subset 4, as well as ABI, ACBD4/5, ANXA13, DIPK2, ELCB, and PCOLCE from subset 5. One additional family, PLSCR, has members in the vicinity of *chst2a* (*plscr1.1*, *plscr1.2*) and *chst2b* (*plscr2*). However, the locations of these genes outside teleost fishes reveal that they likely arose through a local duplication in a bony fish ancestor, at the latest, rather than in teleost fishes and 3R.The identified paralogous segments on chromosomes 2 and 24 in zebrafish, and 17 and 20 in medaka (figure 4B), correspond to chromosome segments previously identified to be the result of 3R (Kasahara et al. 2007; Nakatani et al. 2007; Amores et al. 2011; Nakatani & McLysaght 2017). In Kasahara et al. (2007) and Nakatani et al. (2007) they correspond to pre-3R proto-chromosome “m”, and in Nakatani & McLysaght (2017) they correspond to proto-chromosome “13”. Our analysis also identified the possible location of the *CHST2a* gene in medaka on chromosome 17, had it not been lost early in the euteleost lineage. The chromosome segments for the ABI, ACBD4/5, ANXA13, ELCB and PFK families in the spotted gar, chicken, Western clawed frog, and human genomes (figure 4B) don’t correspond to the *CHST2* and *CHST7*-bearing chromosome segments. However, several studies have identified those chromosome segments to also be part of the *CHST2* and *CHST7*-bearing paralogon (see figure 4 in Kasahara et al. 2007, figure 4 in Bian et al. 2016, and figure 3 in Nakatani & McLysaght 2017).

In the zebrafish, the ABI, NCK, and PFK gene families have members in the vicinity of either *chst2a* or *chst2b* as well as *chst7* (figure 4B), reflecting the 1R/2R-generated paralogy as described above. There are three additional families that also fulfil this criterium, BRINP, NOS1AP, and RGS4/5/8/16 (supplementary figure S6), but are likely not the result of 1R/2R. The single homologous paralogy blocks on human chromosome 1, chicken chromosome 8, Western clawed frog chromosome 4, and spotted gar linkage group 10 suggest that the gene duplicates *BRINP2* and *BRINP3*, as well as *RGS4*, *RGS5*, *RGS8* and *RGS16* arose through ancient local duplications rather than through 1R/2R.

### *CHST4*, *CHST5* and related CHST4/5-like sequences

The branch of the C6OST family that contains the known *CHST4*, *CHST5*, and *CHST6* sequences is by far the most complex of the C6OST phylogeny. This branch of our C6OST phylogeny is shown in figure 5, and the full-species phylogeny is included in supplementary figure S7. Aside from the known C6OST sequences, we can report several *CHST4* and *CHST5*-like subtypes represented in amphibians, non-avian reptiles and jawless vertebrates. The *CHST4* and *CHST5* genes (and *CHST6*) are located on the same chromosome in many species, and some of the *CHST4* and *CHST5*-like genes we identified are located downstream of these, suggesting that there has been a high propensity for the duplication of C6OST genes in these regions.

**Fig. 5.**
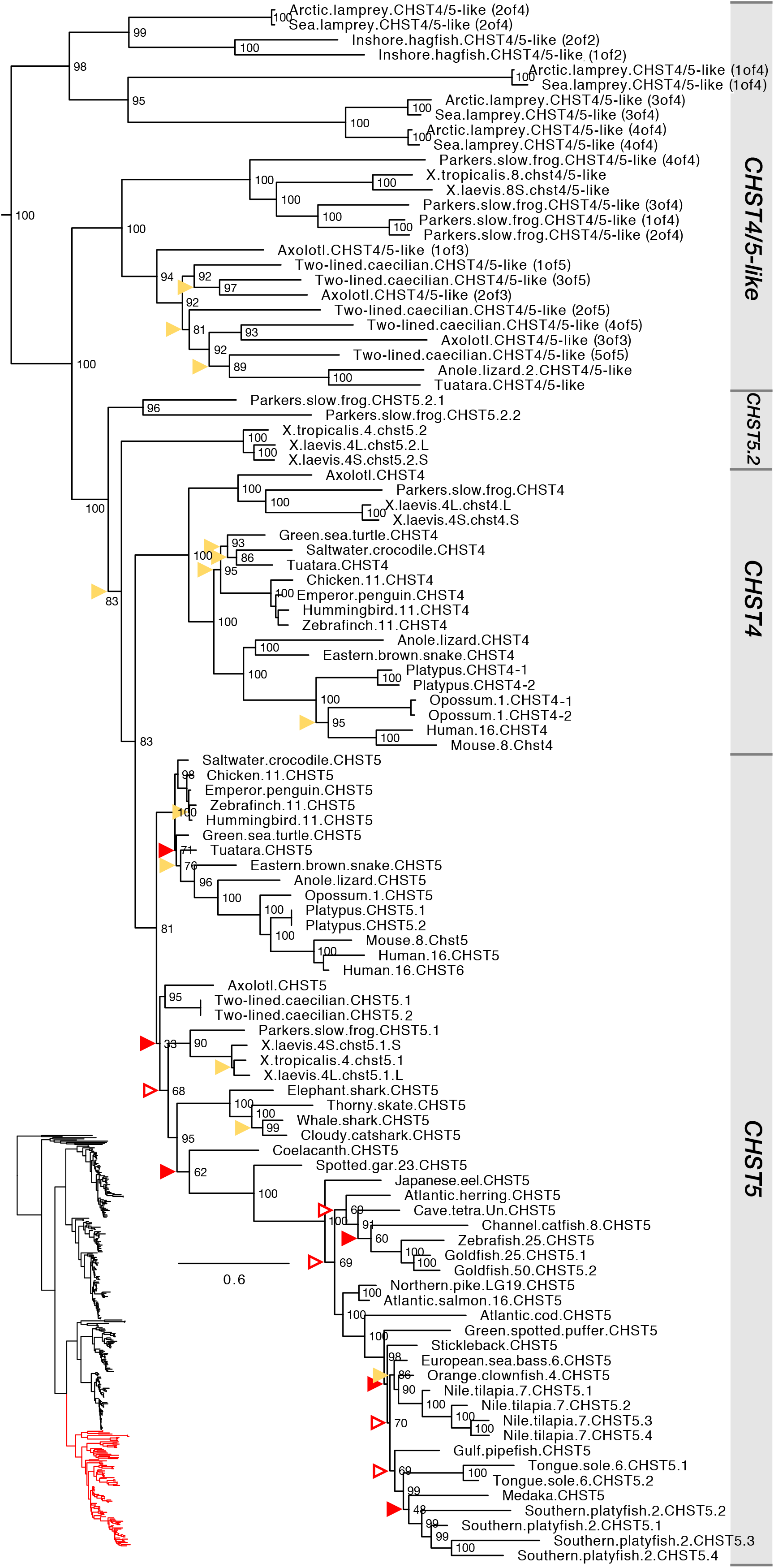
Phylogeny of the *CHST4*, *CHST5* and related C6OST sequences. See figure 2 caption for details.

In both our phylogenies, the known *CHST4* and *CHST5* sequences cluster into two well-defined and well-supported clades. This allowed us to classify a number of sequences with hitherto unclear identities. All identified sequences from cartilaginous fishes, coelacanth, and ray-finned fishes, including spotted gar and teleosts, cluster confidently together with tetrapod sequences within the *CHST5* clade, while *CHST4 sequences* could only be identified from tetrapod species. We could only identify orthologs of the human *CHST6* gene in other primate species, and these cluster confidently within the *CHST5* clade. While there are genes in other mammalian species, chicken and Western clawed frog that have previously been identified as *CHST6* (Taniguchi et al. 2012) our analyses shows that they are better described as *CHST5* for the non-primate mammal and chicken genes, and as *“CHST4*/*5*-like” in the Western clawed frog. The zebrafish *chst5* genes is also erroneously annotated as *chst6* in the zebrafish information network database at zfin.org (gene ID: ZDB-GENE-060810-74). In teleost fishes, we could identify many lineage-specific duplicates of *CHST5*. The largest number was found in the genome of the climbing perch (order *Anabantiformes*) with 15 gene duplicates, some of which are likely pseudogenes. There are also *CHST5* duplicates in the tongue sole (order *Pleuronectiformes*), corkwing wrasse (order *Labriformes*), brown dottyback (order *Pseudochromidae*), turquoise killifish, guppy, and Southern platyfish (order *Cyprinodontiformes*), Eastern happy, zebra mbuna, Nile tilapia, and *Simochromis diagramma* (order *Cichliformes*), European pilchard (order *Clupeiformes*), as well as in the mormyrids, Asian arowana, silver arowana, and Arapaima (superorder *Osteoglossomorpha*) (supplementary figure S7). At least some of the duplications are shared between several species within *Cyprinodontiformes, Cichliformes* and *Osteoglossomorpha.* In the genomes that have been mapped to chromosomes, assembled into linkage groups or longer genomic scaffolds, these duplicated *CHST5* genes are located in tandem rather than on different chromosomes (supplementary data S1). We could also identify *CHST5* duplicates in the goldfish genome, located on chromosomes 25 and 50, that likely arose in the cyprinid fish whole-genome duplication (see figure 2 in Chen et al. 2018). Conversely, no *CHST5* duplicates from the salmoniform-specific whole-genome duplication seem to have been preserved.

In addition to the *CHST4* and *CHST5* (including *CHST6*) sequences, we could identify a multitude of previously unrecognized *CHST4*/*5*-like sequences in jawless vertebrates, amphibians and non-avian reptiles. With some exceptions detailed below, we have used the name *CHST4*/*5*-like for these sequences. In jawless vertebrates, we identified four *CHST4*/*5*-like sequences in the Arctic lamprey and sea lamprey genomes, respectively and two *CHST4*/*5*-like sequences in the inshore hagfish genome. These sequences form a well-supported clade that clusters at the base of the *CHST4*, *CHST5* and *CHST4*/*5-like* branch (figure 5). As for jawed vertebrates, we could identify a well-supported clade of *CHST4*/*5*-like sequences represented in amphibians as well as *Lepidosauria*, excluding snakes. We could identify sequences of this subtype in all investigated amphibian lineages, including the two-lined caecilian (clade *Gymnophiona*), salamanders (order *Urodela*), and frogs (order *Anura*), as well as in the tuatara (order *Rhynchocephalia*), the ocelot gecko (infraorder *Gekkota*), and several lizards, including the European green lizard (clade *Laterata*), the Carolina anole lizard and the central bearded dragon (suborder *Iguania*). This clade clusters basal to the main *CHST4* and *CHST5* branches in both phylogenies. This indicates that these sequences either represent an ancestral jawed vertebrate C6OST subtype that has not been preserved in any other lineages, or more parsimoniously, *CHST4*/*5*-like gene duplicates with a derived mode of sequence evolution that arose in tetrapods before the split between amphibians and amniotes. A conserved synteny analysis of the Carolina anole lizard *CHST4*/*5*-like sequence on chromosome 2 showed no conservation of synteny with other C6OST gene-bearing regions (supplementary figure S8). In amphibians, there have been multiple rounds of local gene duplication within this clade. While the Western and African clawed frogs only have one *CHST4*/*5*-like gene in this clade, Parker’s slow frog has four copies located in tandem on the same genomic scaffold, and the two-lined caecilian has five (supplementary figure S9). The phylogenetic relationships between different duplicates in different species are not resolved in either phylogeny (figure 5, supplementary figure S7). Apart from these *CHST4*/*5*-like sequences, we could identify *CHST5*-like sequences in the three frog genomes. In the phylogeny with full-species representation (supplementary figure S7), these sequences form a well-supported sister clade to the frog *CHST5* sequences. Owing also to their arrangement downstream of the *CHST5* genes in the Western and African clawed frog genomes (supplementary figure S9), we have called them *CHST5.2* and the frog *CHST5* genes *CHST5.1*. The relationship between the frog *CHST5.1* and *CHST5.2* genes is not reproduced in the smaller phylogeny (figure 5). Indeed, the amphibian branch of *CHST5* is unresolved in both phylogenies, likely due to diverging evolutionary rates within the *CHST5* clade. The amphibian *CHST5* (*CHST5.1* in frogs) sequences have had a lower rate of amino acid substitution, while the frog *CHST5.2* sequences seem to have had an accelerated evolutionary rate, as shown by the branch lengths in both phylogenies. The allotetraploid African clawed frog has duplicate *CHST4, CHST5.1* and *CHST5.2* genes, all located on the homeologous chromosome pair 4L and 4S (supplementary figure S9), totaling seven genes within this branch of the C6OST phylogeny.

We could identify only two gene families showing conserved synteny between either *CHST4* or *CHST5* and another C6OST family member: KLF9/13/14/16 and RPGRIP1. These were detected in the zebrafish genome, with members in the vicinity of *chst5* and *chst2a* (KLF9/13/14/16) or *chst2b* (RPGRIP1). However, this conserved synteny relationship is not reproduced in any of the other genomes we investigated, including the medaka (supplementary data S3). Thus, it is likely the result of chromosome rearrangements in the lineage leading to zebrafish. No other patterns of conserved synteny could be detected with respect to the *CHST4*, *CHST5* and related genes. As with the *CHST3* gene, this lack of conserved synteny suggests that the ancestral *CHST4/5* gene did not arise in the two early vertebrate whole-genome duplications 1R and 2R, but rather that it was present together with the ancestral *CHST2/7* and *CHST1/16* genes as well as the *CHST3* gene before 1R.

### Exon/intron structures of vertebrate C6OST genes

As has been reported previously, most mammalian C6OST genes consist of intron-less ORFs (Lee et al. 1999; Bistrup et al. 1999; Kitagawa et al. 2000; Hemmerich, Lee, et al. 2001), with the exception of *CHST3*, whose protein-coding domain is encoded by two exons (Tsutsumi et al. 1998). Here we can report that the protein-coding domain of the *CHST16* genes also consists of two exons. The position of the intron is different from the *CHST3* intron. These exon structures are common to jawed vertebrates and likely represent the exon/intron structures of the ancestral genes. In jawless vertebrates there seem to have been several independent intron insertions that make it difficult to deduce the ancestral conditions. There are several notable exceptions to the common protein-coding exon/intron structures, all found within teleost fishes: *CHST1a* in *Gobiiformes* (gobies) has five exons (independent intron insertions); *CHST1b* in *Acanthomorpha* (spiny-rayed fishes) has five exons (independent intron insertions); *CHST3a* in *Neoteleostei* has three exons, one additional intron insertion downstream of the ancestral jawed vertebrate *CHST3* intron; *CHST3a* in gobies has four exons, one additional intron insertion downstream of ancestral jawed vertebrate and neoteleost introns; *CHST3b* in *Protacanthopterygii* (including Northern pike and salmoniform fishes) has three exons, one additional intron insertion at a different position than neoteleost *CHST3a*; *CHST5* in *Characiformes* (including the Mexican cave tetra and red-bellied piranha) has two exons (independent intron insertion); *CHST7* in *Syngnathiformes* (seahorses, pipefishes) and gobies has two exons, the intron insertion is shared between the two lineages.

### Invertebrate C6OST genes

In addition to the vase tunicate and fruit fly C6OST sequences included in our phylogeny (figure 1), we could identify putative C6OST sequences from the Florida lancelet (*Branchiostoma floridae*), the hemichordate acorn worm (*Saccoglossus kowalevskii*), the purple sea urchin (*Strongylocentrotus purpuratus*) as well as the honey bee (*Apis mellifera*) and the silk moth (*Bombyx mori*) (supplementary data S1). Ultimately, these sequences were not used in our final phylogenies, however it is worth mentioning that there seem to have been extensive lineage-specific expansions of C6OST genes in all but the insect species. As described above, seven unique C6OST sequences have been described from the vase tunicate (Tetsukawa et al. 2010). We could identify 16 unique C6OST sequences in the Florida lancelet, 31 in the acorn worm, and 30 in the purple sea urchin. All full-length sequences contain the characteristic sulfotransferase domain (Pfam PF00685) and have recognizable 5’PSB and 3’PB motifs (supplementary data S2).

## Discussion

### The evolution of vertebrate C6OST genes

We present the first large-scale analysis and phylogenetic classification of vertebrate carbohydrate 6-O sulfotransferase (C6OST) genes. Our analyses are based on genomic and transcriptomic data from a total of 158 species representing all major vertebrate groups, including 18 mammalian species, 33 avian species in 16 orders, 14 non-avian reptile species, six amphibian species, the basal lobe-finned fish coelacanth, the holostean fishes spotted gar and bowfin, 67 teleost fish species in 35 orders, seven cartilaginous fish species, and 3 jawless vertebrate species. This allowed us to identify a previously unrecognized C6OST subtype gene that we have named *CHST16*, which was lost in amniotes, as well as a multitude of lineage-specific duplicates of the known C6OST genes. We could also identify known C6OST genes in lineages where they were previously unrecognized, like *CHST7* in birds and non-avian reptiles, and *CHST1a* and *CHST1b* duplicates in teleost fishes. Our phylogenetic analyses show that the jawed vertebrate C6OST gene family consists of six ancestral clades forming two main branches. The first branch contains *CHST1*, *CHST3* and *CHST16*, and the second branch contains *CHST2*, *CHST7* and *CHST5*. The *CHST5* clade also includes *CHST4* and *CHST6.* We have identified genes within the six ancestral subtypes in the genomes of several cartilaginous fishes, lobe-finned fishes, including the coelacanth and tetrapods, as well as ray-finned fishes, including the spotted gar and teleost fishes (figure 6).

**Fig. 6.**
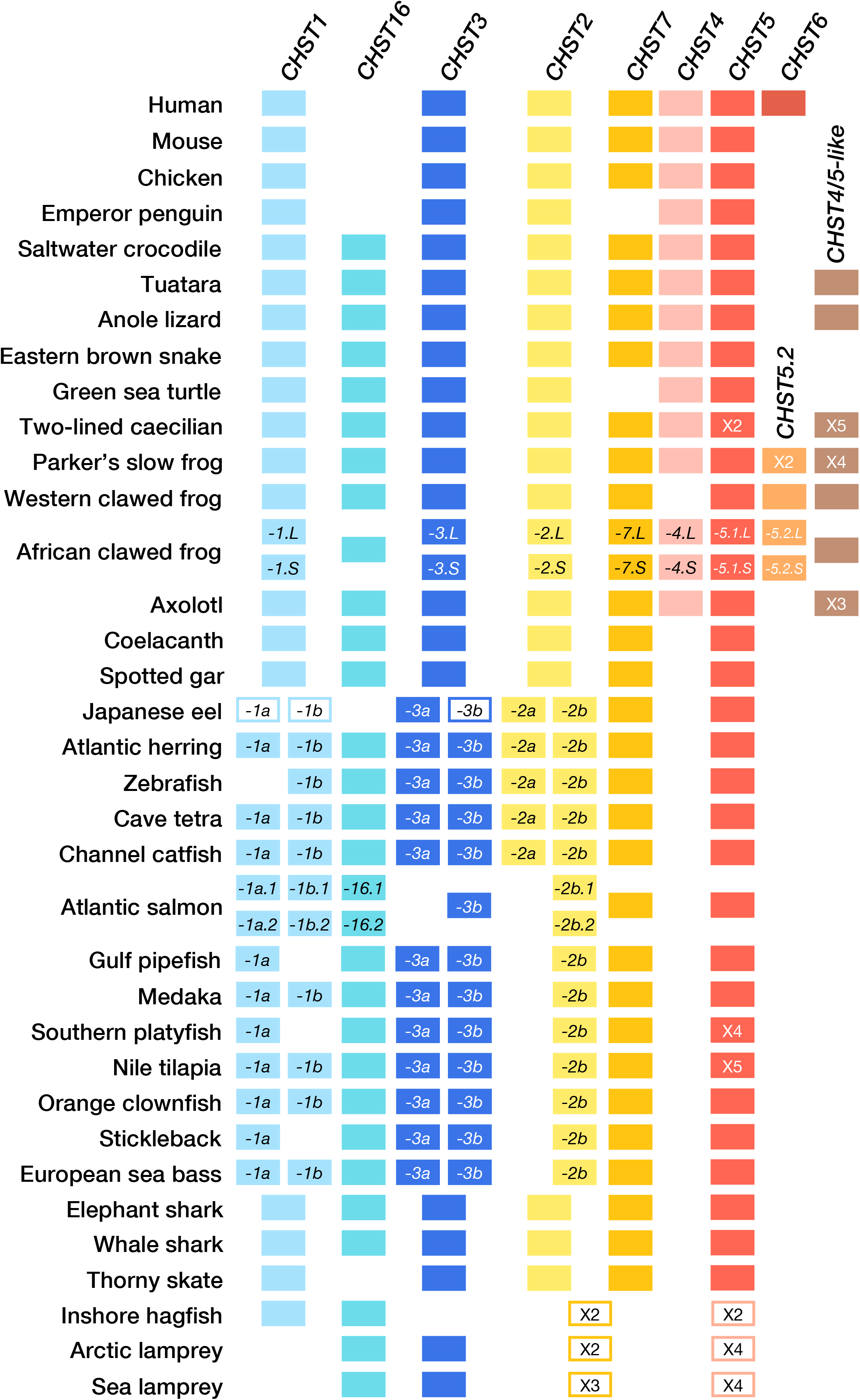
C6OST gene repertoires in selected vertebrate species. Opened boxes indicate genes with unresolved phylogenetic positions. Ancestral teleost, Atlantic salmon and *Xenopus laevis* duplicate gene designations are shown within the boxes. Other species-specific or lineage-specific duplicates are indicated by “X2” etc. within the boxes. Additional *CHST4*/*5-like* and *CHST5.2* genes in non-avian reptiles and amphibians not shown here (see supplementary Fig. S7).

We conducted comparative analyses of conserved synteny - the conservation of gene content across several chromosomal regions, that are the result from block duplications of large chromosomal segments or whole-genome duplications. We found that the basal vertebrate whole-genome duplications (1R/2R) only contributed two additional genes to the ancestral vertebrate repertoire, giving rise to *CHST1* and *CHST16*, as well as *CHST2* and *CHST7* (figure 7). Indeed, the *CHST1* and *CHST16* genes, and the *CHST2* and *CHST7* genes, are located in genomic regions previously recognized to have originated through 1R/2R (Nakatani et al. 2007; Kasahara et al. 2007; Putnam et al. 2008; Sacerdot et al. 2018). Thus, four C6OST genes - *CHST3*, *CHST5* as well as the ancestral *CHST1*/*16* and *CHST2*/*7* - were present already in a vertebrate ancestor, before 1R. Our phylogenetic analysis places the origin of the gene family between the divergence of tunicates from the main chordate branch that took place 518-581 Mya (Hedges and Kumar, 2009) and the emergence of extant vertebrate lineages. Thus, the gene family expansions that laid the ground for the vertebrate C6OST gene family likely occurred very early in chordate/vertebrate evolution.

**Fig. 7.**
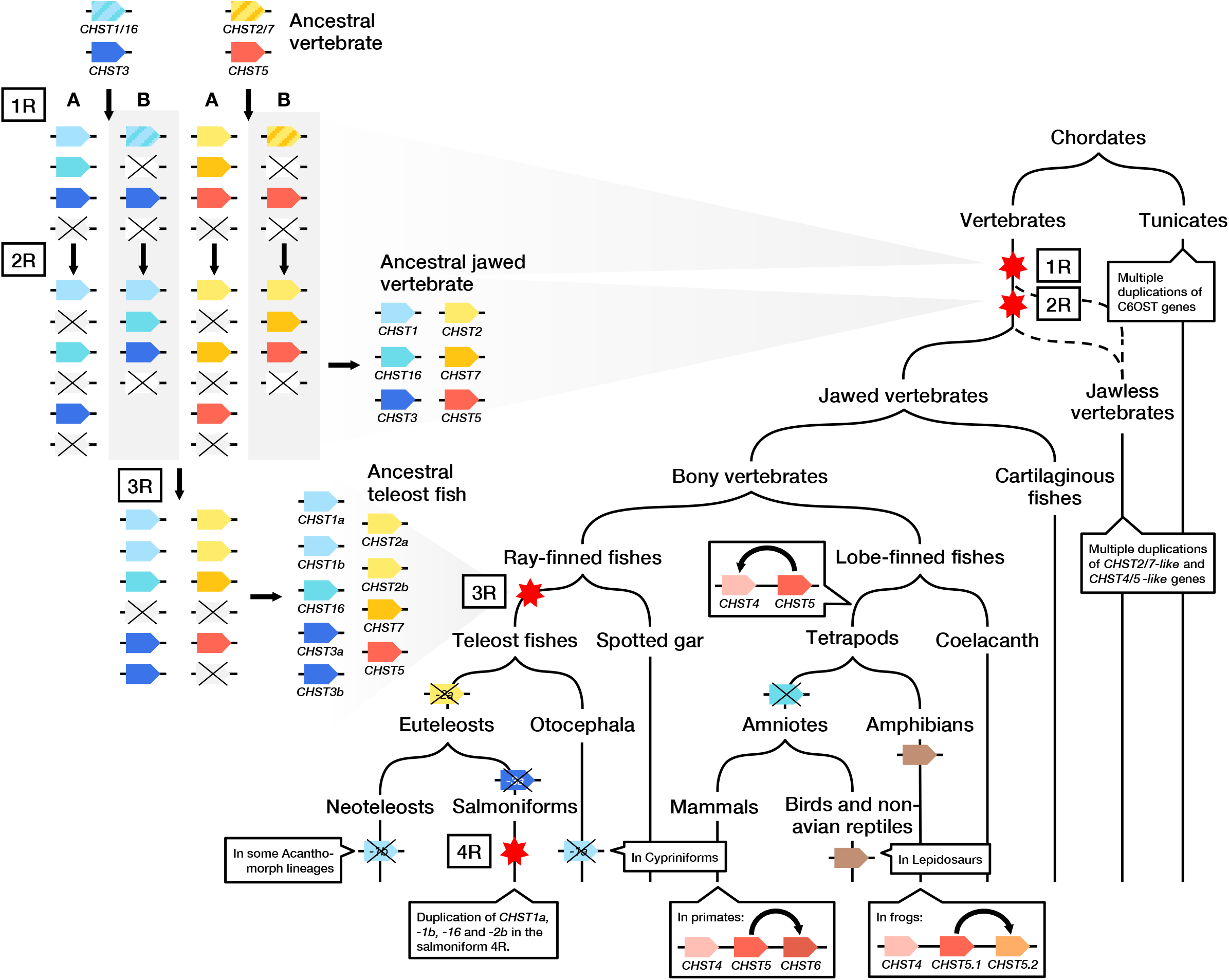
Evolutionary scenario of C6OST family in chordates. Seven-pointed stars indicate whole genome duplications. Crossed-over boxes indicate gene losses. Our analyses indicate that *CHST1* and *CHST16* genes, as well as *CHST2* and *CHST7* genes, arose from ancestral *CHST1*/*16* and *CHST2/7* genes through either 1R (“A”) or 2R (“B”). The uncertain position of jawless vertebrates relative to 1R and 2R is indicated by dashed lines.

In jawless vertebrates (*Agnatha*) we identified the *CHST1* and *CHST16* sequences from the inshore hagfish, as well as the *CHST16* and *CHST3* sequences from the Arctic and sea lampreys (figure 2A and figure 6). We could also identify several *CHST2*/*7*-like, and *CHST4*/*5*-like, sequences in all three jawless vertebrate species, however their duplication events are not completely resolved in our phylogeny (figure 4, figure 5 and open boxes in figure 6). There has been a long debate about whether jawless vertebrates diverged after 1R or whether jawless and jawed vertebrates share both rounds of whole genome duplication (Kuraku et al. 2009). Previous studies of both the sea lamprey and Arctic lamprey genomes indicate that jawless and jawed vertebrates share both whole-genome duplications (Smith et al. 2013; Mehta et al. 2013), but that the shared patterns of gene synteny are obscured by the asymmetric retention and loss of gene duplicates (Kuraku 2013). Another recent reconstruction of vertebrate genome evolution based on the retention of gene content, order and orientation, as well as gene family phylogenies, also concluded that jawless and jawed vertebrate diverged after 2R (Sacerdot et al. 2018; Holland & Ocampo Daza 2018). In contrast, an analysis of the meiotic map of the sea lamprey genome interprets the patterns of synteny conservation differently, suggesting instead one round of genome duplication at the base of vertebrates with subsequent *independent* segmental duplications in jawless and jawed vertebrates (Smith & Keinath 2015). Our results are compatible with at least one whole genome duplication shared between jawless and jawed vertebrates. The *CHST4*/*5*-like genes we identified in jawless vertebrates likely represent the ancestral vertebrate *CHST5* and the *CHST2*/*7*-like genes likely represent the ancestral *CHST2 and CHST7* genes, however, we could not assign them to one individual clade. As our schematic shows both whole-genome duplications were followed by several gene losses (figure 7). Thus, it is also possible that jawless and jawed vertebrates have preserved and lost different gene copies generated by 1R or 2R. Such differential gene losses/preservations or “hidden paralogies” are thought to underlie many ambiguous orthology and paralogy assignments between lamprey sequences and jawed vertebrate sequences (Kuraku 2010).

After the emergence of jawed vertebrates, and later of bony vertebrates, and the split between lobe-finned fishes and ray-finned fishes, the evolution of C6OST genes has taken different routes within different lineages, with several additional gene duplications and losses. The evolution of the *CHST5* branch is particularly marked by local gene duplications: Local gene duplications of *CHST5* generated *CHST4* in a tetrapod ancestor, *CHST6* in a primate ancestor, as well as a previously unrecognized gene in frogs that we called *CHST5.2* (figure 7). Only tetrapods were found to have both *CHST4* and *CHST5*, and all investigated species with assembled genomes have the genes located on the same chromosome, linkage group or genomic scaffold. The cartilaginous fish, coelacanth, spotted gar and teleost sequences within this branch cluster confidently within the *CHST5* clade, rather than basal to the tetrapod *CHST4* and *CHST5* clades. This also indicates that *CHST4* emerged later (figure 7). As detailed above, the phylogenetic identity of the jawless vertebrate *CHST4*/*5*-like sequences is somewhat uncertain, whereby we have decided not to name them *CHST5*. Different genes have previously been identified as *CHST6* in non-primate mammals, chicken, Western clawed frog and zebrafish. In most cases it is *CHST5* that has been misidentified, and we suggest new gene names and symbols in the results above.

The newly identified *CHST16* is present within cartilaginous fishes, but was likely lost from rays and skates. It was also identified in all investigated ray-finned fish species, the coelacanth and amphibians, but not in any of amniote species, indicating an early gene loss in this lineage. Nothing is known about the functions of *CHST16*, so we cannnot speculate whether its deletion was concurrent with a loss of function, or whether another C6OST gene could compensate for its loss. However, it is notable that the likely emergence of *CHST4* as a copy of *CHST5*, and the loss of *CHST16*, are associated with the time window for the transition from water to land and the emergence of the amniotic egg.

The C6OST gene family has had a dynamic and varied evolution within the teleost fish lineage. The teleost-specific whole-genome duplication, 3R, gave rise to duplicates of *CHST1*, *CHST2* and *CHST3*, increasing the number of genes to nine in the teleost ancestor (figure 6, figure 7). This was followed by multiple differential losses; notably, the loss of *CHST1a* from cypriniform fishes, including zebrafish, the loss of *CHST3a* from salmoniform fishes, and of *CHST2a* from euteleost fishes. *CHST1b* seems to have been lost independently within several lineages of spiny-rayed fishes (*Acanthomorpha*). This apparently relaxed selection on the retention of *CHST1b* seems to have been preceded by a stage of rapid evolution. We found that the neoteleost branch of *CHST1b* sequences show an accelerated rate of amino acid substitution coupled with the insertion of four introns into the otherwise uninterrupted ORF. In addition to 3R, there have been more recent whole-genome duplications in cyprinid fishes and in salmoniform fishes (Glasauer & Neuhauss 2014), occasionally called 4R. We concluded that the salmoniform 4R contributed duplicates of *CHST1a*, *CHST1b*, *CHST16* and *CHST2b* (figure 6, figure 7), raising the number of C6OST genes in this lineage to a total of eleven. We identified additional duplicates of *CHST16*, *CHST2a*, *CHST2b*, *CHST3b*, *CHST5* and *CHST7* in the goldfish genome that likely arose in the cyprinid whole-genome duplication, as well as copies of *CHST1b* and *CHST3a* of an uncertain origin, raising the number of C6OST genes in this species to a total of 17. Other notable gene expansions within the teleost fishes include the local expansion of *CHST5* in several lineages, such as killifishes and live-bearers, cichlids and *Osteoglossomorpha*.

Finally, we also identified a number of *CHST4*/*5*-like genes in amphibians and several lineages within *Lepidosauria* whose origins could not be elucidated by phylogenetic nor conserved synteny analyses (figure 7). The species representation suggests that this gene arose early in tetrapod evolution and was subsequently lost multiple times: from mammals, turtles, tortoises and terrapins (*Testudines*), birds and crocodilians (*Archosauria*), as well as snakes. Further investigation of the amphibian *CHST4*/*5*-like sequences and lepidosaurian *CHST4/5*-like sequences in additional amphibian and non-avian reptile species will be needed to resolve this phylogenetic uncertainty.

### Functional considerations

Our interest in the evolution of C6OSTs mainly revolves around their key roles in the formation of extracellular matrices (ECMs) during the formation of mineralized tissues, such as cartilage and bone. Within this context, the modest expansion of the C6OST family in 1R/2R, stands in contrast to several gene families with functions in skeleton formation. Notably the Hox gene family (Ravi et al. 2009; Kuraku & Meyer 2009; Sundström et al. 2008), and the fibrillar collagen gene family (Boot-Handford & Tuckwell 2003; Zhang & Cohn 2008), whose expansions in 1R/2R has been considered essential for the evolution of the vertebrate skeleton (Wada 2010). Several key genes involved in the regulation of bone homeostasis also arose through 1R/2R, including the parathyroid hormone gene (PTH) and the calcitonin genes (CALC), as wells as their cognate receptor genes (Hwang et al. 2013). The bone morphogenic protein (BMP) genes *BMP2* and *BMP4* have also been shown to have emerged in 1R/2R (Marques et al. 2016; Feiner et al. 2019), as have *BMP5*, *BMP6*, *BMP7* and *BMP8*. Sulfotransferases have a pivotal role in generating the active GAGs that are subsequently combined into proteoglycans, building the matrix that provides a scaffold for tissue formation. Our results raise the question of whether enzymes like sulfotransferases were important prerequisites before genes specifically dedicated towards skeleton formation could evolve. Out of the C6OST genes we have studied, *CHST3* and *CHST7* sulfate chondroitin sulfate (CS), and have been implicated in the development and homeostasis of the skeletal system. CS molecules that are 6-O sulfated by *CHST3* are important components of proteoglycans like aggrecan in cartilage. Our results show that *CHST3* and *CHST7* are not closely related, which suggests that chondroitin 6-O sulfation activity has evolved at least twice, even before the advent of cartilaginous skeletons in vertebrates. Indeed, we found that cartilaginous fishes have both *CHST3* and *CHST7* genes, and at least *CHST3* is clearly present in jawless vertebrates. It is also notable that *CHST7* is most closely related to *CHST2*, which suggests some functional overlap, at least in early vertebrate evolution. *CHST2*, through the 6-O sulfation of keratan sulfate (KS), may also have a role in the skeletal system, in the maintenance of cartilaginous tissue (Hayashi et al. 2011). The expression profiles of the zebrafish *chst3a*, *chst3b* and *chst7* genes during development mirror the known functions of *CHST3* and *CHST7* in mammals, with expression mainly in skeletal tissues and the central nervous system (Habicher et al. 2015), albeit with only a very weak signal for *chst3b*. Further study in teleost species should be done in order to compare the expression profiles of the teleost-specific CHST3 duplicates.

As for the novel C6OST subtype gene we describe here, *CHST16*, its close phylogenetic relationship to the *CHST1* paralogue could indicated some similarities in gene expression profile and preferred substrate. A large-scale high-throughput expression analysis identified expression of *chst1b* in the zebrafish retina and brain (Thisse & Thisse 2004), which is shared with mammalian *CHST1* orthologues. In the zebrafish retina, as in other vertebrate retinas, KS is a common and abundant GAG molecule (Souza et al. 2007). These facts point at a common role of CHST1 in 6-O sulfation of KS in the retina and central nervous system across lobe-finned fishes, including tetrapods and ray-finned fishes, including teleost fishes.

## Supporting information

Supplementary Dara S2

## Acknowledgments

This work was supported by the Swedish Research Council (Vetenskapsrådet) through an International Postdoc Grant awarded to D.O.D. (2016-00552) and a Starting Grant awarded to T.H. (621-2012-4673). We wish to thank the G10K Consortium and professor Erich D. Jarvis from Rockefeller University, New York, for access to genomic data from several key species, including platypus, greater horseshoe bat, Anna’s hummingbird, kakapo, zebra finch, two-lined caecilian, and thorny skate, through the Vertebrate Genomes Project (vertebrategenomesproject.org).

