## Supplementary Table S1-S2, Supplementary Fig. S1-S9 for "Reconstruction of the carbohydrate 6-O sulfotransferase gene family evolution in vertebrates reveals novel member, CHST16, lost in amniotes"

**Supplementary Material**

**Supplementary Table S1.** All investigated species and genome/transcriptome assemblies.

**Supplementary Table S2.** Neighboring gene families around C6OST genes.

**Supplementary Fig. S1.** Maximum likelihood phylogeny of *CHST1* sequences with full species representation. The phylogeny is supported by approximate Likelihood-Ratio Test (aLRT) and UltraFast Bootstrap (UFBoot) analyses, and rooted with the inshore hagfish *CHST1* sequence. Both aLRT and UFBoot supports are shown, in that order. aLRT supports have been rounded to the nearest integer. Sequence names include species name abbreviations (see Supplementary table 1) followed by chromosome/linkage group designations (if available) and gene symbols. Asterisks indicate incomplete sequences. The fast-evolving branch of neoteleost *CHST1b* sequences is circled.

**Supplementary Fig. S2.** Maximum likelihood phylogeny of *CHST16* sequences with full species representation. The phylogeny is rooted with the cyclostome *CHST16* sequences. See Fig. S1 caption for more details.

**Supplementary Fig. S3.** Maximum likelihood phylogeny of *CHST3* sequences with full species representation. The phylogeny is rooted with the lamprey *CHST3* sequences. See Fig. S1 caption for more details.

**Supplementary Fig. S4.** Maximum likelihood phylogeny of *CHST2* sequences with full species representation. The phylogeny is rooted with the cartilaginous fish *CHST2* sequences due to the ambiguous identity of the cyclostome *CHST2/7*-like sequences. See Fig. S1 caption for more details.

**Supplementary Fig. S5.** Maximum likelihood phylogeny of *CHST2* sequences with full species representation. The phylogeny is rooted with the cartilaginous fish *CHST7* sequences due to the ambiguous identity of the cyclostome *CHST2/7*-like sequences. See Fig. S1 caption for more details.

**Supplementary Fig. S6.** Alternative conserved synteny between CHST2 and CHST7-bearing chromosome regions in tetrapods and the spotted gar, including CHST2a and CHST2b-bearing regions in teleost fishes.. Stars indicate chromosome regions in common with analysis on figure 4B.

**Supplementary Fig. S7.** Maximum likelihood phylogeny of *CHST4, CHST5* and related sequences with full species representation. The phylogeny is rooted with the non-avian reptile and amphibian "*CHST4/*5-like" sequences. See Fig. S1 caption for more details.

**Supplementary Fig. S8.** Conserved synteny between the anole lizard "*CHST4/5-like*" gene-bearing region on chromosome 2 and the corresponding human and spotted gar chromosome regions. The gene predictions located within a segment of 5 MB in each direction of the *CHST4/5-like* gene were identified as described in Methods, and the orthologous gene locations in the human and spotted gar genomes were identified in the Ensembl database.

**Supplementary Fig. S9.** "*CHST4/5*-like" genes in amphibians and non-avian reptiles. Chromosome positions are given in megabases (Mb).

**Supplementary Data S1.** Identified C6OST sequence data. Includes chromosomal locations and database identifiers of all C6OST sequences identified in this study.

**Supplementary Data S2.** All C6OST sequences identified in this study in alphabetical order by species. Sequence names include species abbreviations (Supplementary table 1) followed by chromosome/linkage group designations (if available) and gene symbols. Asterisks indicate incomplete sequences. Numbers after / indicate sequence length. Also includes a list of identical sequences.

**Supplementary Data S3.** Neighboring gene family data. Includes chromosomal locations and database identifiers of all neighboring genes in the human, chicken Western clawed frog, spotted gar, zebrafish, medaka, elephant shark, vase tunicate and fruit fly. Also includes all identified synteny blocks.

**Supplementary Table S1.** All investigated species and genome/transcriptome assemblies.

| | Order | Species | Common name | Abbr. | Data type | Chr./LG | Assembly version | Database | Link |
|---|---|---|---|---|---|---|---|---|---|
| Mammals | *Monotremata* | *Ornitorhynchus anatinus* | Platypus | Oan | Genome | No | Ornithorhynchus_anatinus-5.0.1 | NCBI | [LINK] |
| | | *Ornitorhynchus anatinus* | Platypus | Oan | Genome | No | mOrnAna1_t3.p | VGP | [LINK] |
| | *Didelphimorphia* | *Monodelphis domestica* | Gray short-tailed opossum | Mdo | Genome | Yes | MonDom5 | NCBI | [LINK] |
| | *Diprotodontia* | *Phascolarctos cinereus* | Koala | Pci | Genome | No | phaCin_unsw_v4.1 | NCBI | [LINK] |
| | *Primates* | *Homo sapiens* | Human | Hsa | Genome | Yes | GRCh38.p12 | NCBI | [LINK] |
| | | *Pan troglodytes* | Chimpanzee | Ptr | Genome | Yes | Clint_PTRv2 | NCBI | [LINK] |
| | | *Macaca mulatta* | Rhesus macaque | Mml | Genome | Yes | Mmul_8.0.1 | NCBI | [LINK] |
| | | *Saimiri boliviensis boliviensis* | Bolivian squirrel monkey | Sbo | Genome | No | SaiBol1.0 | NCBI | [LINK] |
| | | *Callithrix jacchus* | Marmoset | Cja | Genome | Yes | Callithrix jacchus-3.2 | NCBI | [LINK] |
| | *Rodentia* | *Mus musculus* | Mouse | Mmu | Genome | Yes | GRCm38.p6 | NCBI | [LINK] |
| | *Lagomorpha* | *Oryctolagus cuniculus* | Rabbit | Oca | Genome | Yes | OryCun2.0 | NCBI | [LINK] |
| | *Eulipotyphla* | *Condylura cristata* | Star-nosed mole | Ccr | Genome | No | ConCri1.0 | NCBI | [LINK] |
| | *Chiroptera* | *Desmodus rotundus* | Common vampire bat | Dro | Genome | No | ASM294091v2 | NCBI | [LINK] |
| | | *Rhinolophus ferrumequinum* | Greater horseshoe bat | Rfe | Genome | No | mRhiFer1_v1 | VGP | [LINK] |
| | *Cetartiodactyla* | *Physeter catodon* | Sperm whale | Pca | Genome | Yes | PhyCat2.0 | NCBI | [LINK] |
| | | *Sus scrofa* | Pig | Ssc | Genome | Yes | Sscrofa11.1 | NCBI | [LINK] |
| | *Carnivora* | *Canis lupus familiaris* | Domestic dog | Cfa | Genome | Yes | CanFam3.1 | NCBI | [LINK] |
| | *Proboscidea* | *Loxodonta africana* | African savanna elephant | Laf | Genome | No | Loxafr3.0 | NCBI | [LINK] |
| | *Cingulata* | *Dasypus novemcinctus* | Nine-banded armadillo | Dno | Genome | No | Dasnov3.0 | NCBI | [LINK] |
| Birds | *Rheiformes* | *Rhea americana* | Greater rhea | Ram | Genome | No | rheAme1 | NCBI | [LINK] |
| | *Galliformes* | *Numida meleagris* | Helmeted guineafowl | Nme | Genome | Yes | NumMel1.0 | NCBI | [LINK] |

| | | | | | | | | |
|---|---|---|---|---|---|---|---|---|
| | *Gallus gallus* | Chicken | Gga | Genome | Yes | GRCg6a | NCBI | [LINK] |
| | *Tympanuchus cupido pinnatus* | Greater prairie chicken | Tcu | Genome | No | T_cupido_pinnatus_GPC_3440_v1 | NCBI | [LINK] |
| | *Meleagris gallopavo* | Turkey | Mga | Genome | Yes | Turkey_5.0 | NCBI | [LINK] |
| *Anseriformes* | *Anas platyrhynchos* | Mallard duck | Apl | Genome | No | BGI_duck_1.0 | NCBI | [LINK] |
| | *Anas platyrhynchos* | Mallard duck | Apl | Genome | Yes | IASCAAS_PekingDuck_PBH1.5 | NCBI | [LINK] |
| | *Anser cygnoides* | Swan goose | Acy | Genome | No | GooseV1.0 | NCBI | [LINK] |
| | *Anser brachyrhynchus* | Pink-footed goose | Abr | Genome | No | ASM259213v1 | NCBI | [LINK] |
| *Apodiformes* | *Calypte anna* | Anna's hummingbird | Can | Genome | Yes | bCalAnn1_v1 | VGP | [LINK] |
| *Cuculiformes* | *Cuculus canorus* | Common cuckoo | Ccn | Genome | No | ASM70932v1 | NCBI | [LINK] |
| *Columbiformes* | *Columba livia* | Rock pigeon | Cli | Genome | Yes | colLiv2 | NCBI | [LINK] |
| | *Patagioenas fasciata monilis* | Band-tailed pigeon | Pfa | Genome | No | NIATT_ARIZONA | NCBI | [LINK] |
| *Charadriiformes* | *Uria lomvia* | Thick-billed murre/guillemot | Ulo | Genome | No | UriLom_1.1 | NCBI | [LINK] |
| *Sphenisciformes* | *Aptenodytes forsteri* | Emperor penguin | Afo | Genome | No | ASM69914v1 | NCBI | [LINK] |
| | *Spheniscus mendiculus* | Galápagos penguin | Sme | Genome | No | ASM326465v1 | NCBI | [LINK] |
| | *Pygoscelis antarcticus* | Chinstrap penguin | Pan | Genome | No | ASM326459v1 | NCBI | [LINK] |
| *Suliformes* | *Nannopterum harrisi* | Galápagos flightless cormorant | Nha | Genome | No | Pharrisi_ref_V1 | NCBI | [LINK] |
| *Ophistocomiformes* | *Ophistocomus hoazin* | Hoatzin | Oho | Genome | No | ASM69207v1 | NCBI | [LINK] |
| *Accipitriformes* | *Aquila chrysaetos chrysaetos* | European golden eagle | Ach | Genome | No | bAquChr1.1 | NCBI | [LINK] |
| *Strigiformes* | *Strix occidentalis caurina* | Spotted owl | Soc | Genome | No | Soccid_v01 | NCBI | [LINK] |
| *Piciformes* | *Picoides pubescens* | Downy woodpecker | Ppu | Genome | No | ASM69900v1 | NCBI | [LINK] |
| *Falconiformes* | *Falco peregrinus* | Peregrine falcon | Fpe | Genome | No | F_peregrinus_v1.0 | NCBI | [LINK] |
| | *Falco cherrug* | Saker falcon | Fch | Genome | No | F_cherrug_v1.0 | NCBI | [LINK] |
| *Psittaciformes* | *Melopsittacus undulatus* | Budgerigar | Mun | Genome | No | Melopsittacus_undulatus_6.3 | NCBI | [LINK] |
| | *Strigops habroptilus* | Kakapo | Sha | Genome | No | bStrHab1_v1 | VGP | [LINK] |

| | | | | | | | | | |
|---|---|---|---|---|---|---|---|---|---|
| | *Passeriformes* | *Lepidothrix coronata* | Blue-crowned manakin | Lco | Genome | No | Lepidothrix_coronata-1.0 | NCBI | [LINK] |
| | | *Corvus cornix cornix* | Hooded crow | Cco | Genome | No | ASM73873v2 | NCBI | [LINK] |
| | | *Parus major* | Great tit | Pmj | Genome | Yes | Parus_major1.1 | NCBI | [LINK] |
| | | *Ficedula albicollis* | Collared flycatcher | Fal | Genome | Yes | FicAlb1.5 | NCBI | [LINK] |
| | | *Taeniopygia guttata* | Zebra finch | Tgu | Genome | Yes | Taeniopygia_guttata-3.2.4 | NCBI | [LINK] |
| | | *Taeniopygia guttata* | Zebra finch | Tgu | Genome | No | bTaeGut1_v1 | VGP | [LINK] |
| | | *Passer domesticus* | House sparrow | Pdo | Genome | Yes | Passer_domesticus-1.0 | NCBI | [LINK] |
| Non-avian reptiles | *Crocodylia* | *Alligator mississipiensis* | American alligator | Ami | Genome | No | ASM28112v4 | NCBI | [LINK] |
| | | *Crocodylus porosus* | Australian saltwater crocodile | Cpo | Genome | No | CroPor_comp1 | NCBI | [LINK] |
| | *Testudines* | *Chrysemys picta bellii* | Western painted turtle | Cpi | Genome | Yes | Chrysemys_picta_bellii-3.0.3 | NCBI | [LINK] |
| | | *Chelonia mydas* | Green sea turtle | Cmy | Genome | No | CheMyd_1.0 | NCBI | [LINK] |
| | | *Pelodiscus sinensis* | Chinese soft-shell turtle | Psi | Genome | No | PelSin_1.0 | NCBI | [LINK] |
| | *Rhynchocephalia* | *Sphenodon punctatus* | Tuatara | Spp | Genome | No | ASM311381v1 | NCBI | [LINK] |
| | *Gekkota* | *Paroedura picta* | Ocelot gecko | Ppi | Genome | No | Ppicta_assembly_v1 | NCBI | [LINK] |
| | *Laterata* | *Lacerta viridis* | European green lizard | Lvi | Genome | No | ASM90024590v1 | NCBI | [LINK] |
| | *Iguania* | *Anolis carolinensis* | Carolina anole lizard | Aca | Genome | Yes | AnoCar2.0 | NCBI | [LINK] |
| | | *Pogona vitticeps* | Central bearded dragon | Pvi | Genome | No | pvi1.1 | NCBI | [LINK] |
| | *Serpentes* | *Protobothrops flavoviridis* | Habu (Ryukyu Islands pit viper) | Pfl | Genome | No | HabAm_1.0 | NCBI | [LINK] |
| | | *Thamnophis sirtalis* | Common garter snake | Tsi | Genome | No | Thamnophis_sirtalis-6.0 | NCBI | [LINK] |
| | | *Pseudonaja texilis* | Eastern brown snake | Pte | Genome | No | EBS10Xv2-PRI | NCBI | [LINK] |
| | | *Python bivittatus* | Burmese python | Pbi | Genome | No | Python_molurus_bivittatus-5.0.2 | NCBI | [LINK] |
| Amphibians | *Gymnophiona* | *Rhinatrema bivittatum* | Two-lined caecilian | Rbi | Genome | No | aRhiBiv1_t4.p | VGP | [LINK] |

| | | | | | | | | | |
|---|---|---|---|---|---|---|---|---|---|
| | *Caudata* | *Ambystoma mexicanum* | Axolotl | Ame | Genome | No | AxolotlMexicanum_assembly.v1 | Ambystoma.org | [LINK] |
| | | *Ambystoma mexicanum* | Axolotl | Ame | Genome | No | ASM291563v2 | NCBI | [LINK] |
| | | *Notophthalmus viridescens* | Eastern newt | Nvi | Transcriptome | N/A | Feb 2014 | Sandberg lab | [LINK] |
| | *Anura* | *Nanorana parkeri* | Parker's slow frog | Npa | Genome | No | ASM93562v1 | NCBI | [LINK] |
| | | *Xenopus (Silurana) tropicalis* | Western clawed frog | Xtr | Genome | Yes | JGI v9.1 | NCBI | [LINK] |
| | | *Xenopus laevis* | African clawed frog | Xla | Genome | Yes | JGI v9.2/ Xenopus_laevis_v2 | NCBI | [LINK] |
| Lobe-finned fishes | *Actinistia* | *Latimeria chalumnae* | Coelacanth | Lch | Genome | No | LatCha1 | NCBI | [LINK] |
| Ray-finned fishes | *Holostei* | *Amia calva* | Bowfin | Acl | Transcriptome | N/A | N/A | PhyloFish | [LINK] |
| | | *Lepisosteus oculatus* | Spotted gar | Loc | Genome | Yes | LepOcu1 | NCBI | [LINK] |
| Teleosts | *Osteoglossomorpha* | *Pantodon buchholzi* | African butterflyfish | Pbu | Transcriptome | N/A | N/A | PhyloFish | [LINK] |
| | | *Gnathonemus petersii* | Peters' elephantnose fish | Gpe | Transcriptome | N/A | N/A | PhyloFish | [LINK] |
| | | *Paramormyrops kingsleyae* | Old Calabar mormyrid | Pki | Genome | No | PKINGS_0.1 | NCBI | [LINK] |
| | | *Scleropages formosus* | Asian arowana (golden) | Sfo | Genome | No | ASM162426v1 | NCBI | [LINK] |
| | | *Osteoglossum bicirrhosum* | Silver arowana | Obi | Transcriptome | N/A | N/A | PhyloFish | [LINK] |
| | | *Arapaima gigas* | Arapaima | Agi | Genome | No | GCA_900497675.1 | NCBI | [LINK] |
| | *Elopomorpha* | *Anguilla anguilla* | European eel | Aan | Transcriptome | N/A | N/A | PhyloFish | [LINK] |
| | | *Anguilla anguilla* | European eel | Aja | Genome | No | Anguilla_anguilla_v1_09_nov_10 | NCBI | [LINK] |
| | | *Anguilla japonica* | Japanese eel | Aja | Genome | No | Ajaponica_ver_D2 | NCBI | [LINK] |
| | *Clupeiformes* | *Clupea harengus* | Atlantic herring | Cha | Genome | No | ASM96633v1 | NCBI | [LINK] |
| | | *Alosa alosa* | Allis shad | Aal | Transcriptome | N/A | N/A | PhyloFish | [LINK] |
| | | *Sardina pilchardus* | European pilchard | Spi | Genome | No | SP_G | NCBI | [LINK] |
| | *Cypriniformes* | *Danio rerio* | Zebrafish | Dre | Genome | Yes | GRCz11 | NCBI | [LINK] |
| | | *Danionella dracula* | Dracula fish | Ddr | Genome | No | fDanDra1.1 | NCBI | [LINK] |

| | | | | | | | | |
|---|---|---|---|---|---|---|---|---|
| | *Leuciscus waleckii* | Amur ide | Lwa | Genome | Yes | GCA_900092035.1 | NCBI | [LINK] |
| | *Cyprinus carpio* | Common carp | Cca | Genome | Yes | GCA_000951615.2 | NCBI | [LINK] |
| | *Carassius auratus* | Goldfish | Cau | Genome | Yes | ASM336829v1 | NCBI | [LINK] |
| *Characiformes* | *Astyanax mexicanus* | Mexican cave tetra | Amx | Genome | Yes | Astyanax_mexicanus-2.0 | NCBI | [LINK] |
| | *Pygocentrus nattereri* | Red-bellied piranha | Pna | Genome | No | Pygocentrus_nattereri-1.0.2 | NCBI | [LINK] |
| *Gymnotiformes* | *Electrophorus electricus* | Electric eel | Eel | Genome | No | Ee_SOAP_WITH_SSPACE | NCBI | [LINK] |
| *Siluriformes* | *Ictalurus punctatus* | Channel catfish | Ipu | Genome | Yes | IpCoco_1.2 | NCBI | [LINK] |
| *Esociformes* | *Esox lucius* | Northern pike | Elu | Genome | Yes | Eluc_V3 | NCBI | [LINK] |
| *Salmoniformes* | *Salmo salar* | Atlantic salmon | Ssa | Genome | Yes | ICSASG_v2 | NCBI | [LINK] |
| | *Salvelinus alpinus* | Arctic char | Sal | Genome | Yes | ASM291031v2 | NCBI | [LINK] |
| | *Oncorhynchus mykiss* | Rainbow trout | Omy | Genome | Yes | Omyk_1.0 | NCBI | [LINK] |
| *Osmeriformes* | *Osmerus eperlanus* | European smelt | Oep | Genome | No | ASM90030227v1 | NCBI | [LINK] |
| *Lampriformes* | *Lampris guttatus* | Opah | Lgu | Genome | No | ASM90030254v1 | NCBI | [LINK] |
| | *Regalecus glesne* | Giant oarfish | Rgl | Genome | No | ASM90030258v1 | NCBI | [LINK] |
| *Gadiformes* | *Gadus morhua* | Atlantic cod | Gmo | Genome | No | GadMor_May2010 | NCBI | [LINK] |
| *Beryciformes* | *Monocentris japonica* | Pineconefish | Mja | Genome | No | ASM90032336v1 | NCBI | [LINK] |
| *Holocentriformes* | *Holocentrus rufus* | Longspine squirrefish | Hru | Genome | No | ASM90030261v1 | NCBI | [LINK] |
| | *Myripristis jacobus* | Blackbar soldierfish | Mjc | Genome | No | ASM90030255v1 | NCBI | [LINK] |
| *Ophidiiformes* | *Brotula barbata* | Bearded brotula | Bba | Genome | No | ASM90030326v1 | NCBI | [LINK] |
| *Gobiiformes* | *Boleophthalmus pectinirostris* | Great bluespotted mudskipper | Bpe | Genome | No | BP.fa | NCBI | [LINK] |
| | *Periophthalmus magnuspinnatus* | N/D | Pmg | Genome | No | PM.fa | NCBI | [LINK] |
| *Scombriformes* | *Thunnus albacares* | Yellowfin tuna | Tal | Genome | No | ASM90030262v1 | NCBI | [LINK] |
| *Syngnathiformes* | *Hippocampus comes* | Tiger tail seahorse | Hco | Genome | No | H_comes_QL1_v1 | NCBI | [LINK] |
| | *Syngnathus scovelli* | Gulf pipefish | Sco | Genome | Yes | ssc_2016_12_20_chromlevel | Cresko lab | [LINK] |

| | | | | | | | | |
|---|---|---|---|---|---|---|---|---|
| *Synbranchiformes* | *Mastacembelus armatus* | Zig-zag eel/tire-track eel | Mar | Genome | No | fMasArm1.1 | NCBI | [LINK] |
| *Anabantiformes* | *Anabas testudineus* | Climbing perch | Ate | Genome | No | fAnaTes1.1 | NCBI | [LINK] |
| *Carangiformes* | *Seriola quinqueradiata* | Japanese amberjack | Squ | Genome | No | Squ_2.0 | NCBI | [LINK] |
| *Centropomidae* | *Lates calcarifer* | Barramundi | Lcl | Genome | No | ASM164080v1 | NCBI | [LINK] |
| *Pleuronectiformes* | *Scophthalmus maximus* | Turbot | Sma | Genome | Yes | ASM318616v1 | NCBI | [LINK] |
| | *Cynoglossus semilaevis* | Tongue sole | Cse | Genome | Yes | Cse_v1.0 | NCBI | [LINK] |
| *Beloniformes* | *Oryzias latipes* | Medaka | Ola | Genome | Yes | ASM223467v1 | NCBI | [LINK] |
| *Cyprinodontiformes* | *Nothobranchius furzeri* | Turquoise killifish | Nfu | Genome | Yes | Nfu_20140520 | NCBI | [LINK] |
| | *Poecilia reticulata* | Guppy | Pre | Genome | No | Guppy_female_1.0+MT | NCBI | [LINK] |
| | *Xiphophorus maculatus* | Southern platyfish | Xma | Genome | Yes | X_maculatus-5.0-male | NCBI | [LINK] |
| *Cichliformes* | *Astatotilapia calliptera* | Eastern happy | Asc | Genome | Yes | fAstCal1.2 | NCBI | [LINK] |
| | *Maylandia zebra* | Zebra mbuna | Mze | Genome | Yes | M_zebra_UMD2a | NCBI | [LINK] |
| | *Oreochromis niloticus* | Nile tilapia | Oni | Genome | Yes | Orenil1.1 | NCBI | [LINK] |
| | *Simochromis diagramma* | N/D | Sdi | Genome | No | fSimDia1.1 | NCBI | [LINK] |
| | *Amphilophus citrinellus* | Midas cichlid | Aci | Genome | No | Midas_v5 | NCBI | [LINK] |
| *Pomacentridae* | *Amphiprion percula* | Orange clownfish | Ape | Genome | Yes | Nemo_v1.1 | NCBI | [LINK] |
| *Pseudochromidae* | *Pseudochromis fuscus* | Brown dottyback | Pfu | Genome | No | ASM90032334v1 | NCBI | [LINK] |
| *Labriformes* | *Symphodus melops* | Corkfin wrasse | Sml | Genome | No | ASM281910v1 | NCBI | [LINK] |
| *Centrarchiformes* | *Oplegnathus fasciatus* | Barred knifejaw | Ofa | Genome | No | ASM341684v1 | NCBI | [LINK] |
| | *Maccullochella peelii* | Murray cod | Mpe | Genome | No | mcod_v1 | NCBI | [LINK] |
| *Perciformes* | *Perca fluviatilis* | European perch | Pfv | Genome | No | UTU_Pfluv_1.1 | NCBI | [LINK] |
| | *Sebastes nigrocinctus* | Tiger rockfish | Sni | Genome | No | ASM47523v3 | NCBI | [LINK] |
| | *Gasterosteus aculeatus* | Three-spined stickleback | Gac | Genome | Yes | BROAD S1 | Ensembl | [LINK] |
| *Moroniformes* | *Dicentrarchus labrax* | European sea bass | Dla | Genome | No | seabass_V1.0 | NCBI | [LINK] |
| *Sciaeniformes* | *Larimichthys crocea* | Large yellow croaker | Lcr | Genome | Yes | L_crocea_2.0 | NCBI | [LINK] |
| *Spariformes* | *Sparus aurata* | Gilthead seabream | Sau | Genome | Yes | ASM330901v1 | NCBI | [LINK] |

| | | | | | | | | | |
|---|---|---|---|---|---|---|---|---|---|
| | | *Spondyliosoma cantharus* | Black seabream | Sca | Genome | No | ASM90030268v1 | NCBI | [LINK] |
| | *Tetraodontiformes* | *Mola mola* | Ocean sunfish | Mmo | Genome | No | ASM169857v1 | NCBI | [LINK] |
| | | *Takifugu rubripes* | Japanese pufferfish (fugu) | Tru | Genome | Yes | FUGU5 | NCBI | [LINK] |
| | | *Tetraodon nigroviridis* | Green spotted pufferfish | Tni | Genome | Yes | TETRAODON 8.0 | Ensembl | [LINK] |
| Cartilaginous fishes | *Holocephala* | *Callorhinchus milii* | Elephant shark | Cmi | Genome | No | Callorhinchus_milii-6.1.3 | NCBI | [LINK] |
| | *Batoidea* | *Amblyraja radiata* | Thorny skate | Ara | Genome | No | sAmbRad1_p1 | VGP | [LINK] |
| | | *Leucoraja erinacea* | Little skate | Ler | Transcriptome | N/A | GSE93582 | NCBI | [LINK] |
| | | *Okamejei kenojei* | Ocellate spot skate | Oke | Transcriptome | N/A | Data2Okenojeitrinity | Figshare | [LINK] |
| | *Selachimorpha* | *Chiloscyllium punctatum* | Brownbanded bambooshark | Cpu | Genome | No | Cpunctatum_v1.0 | NCBI | [LINK] |
| | | *Rhincodon typus* | Whale shark | Rty | Genome | No | ASM164234v2 | NCBI | [LINK] |
| | | *Rhincodon typus* | Whale shark | Rty | Genome | No | Rtypus_kobe_v1.0 | Figshare | [LINK] |
| | | *Scyliorhinus torazame* | Cloudy catshark | Sto | Genome | No | Storazame_v1.0 | NCBI | [LINK] |
| Cyclostomes | *Myxiniformes* | *Eptatretus burgeri* | Inshore hagfish | Ebu | Genome | No | Eburgeri_3.2 | Ensembl | [LINK] |
| | *Petromyzontiformes* | *Lethenteron camtschaticum* | Arctic (Japanese) lamprey | Lca | Genome | No | LetJap1.0 | NCBI | [LINK] |
| | | *Petromyzon marinus* | Sea lamprey | Pma | Genome | No | Pmar_germline 1.0 | NCBI | [LINK] |
| Tunicates | *Tunicata* | *Ciona intestinalis* | Vase tunicate | Cin | Genome | Yes | KH | NCBI | [LINK] |
| Cephalochordates | *Cephalochordata* | *Branchiostoma floridae* | Florida lancelet | Bfl | Genome | No | Version 2 | NCBI | [LINK] |
| Hemichordates | *Enteropneusta* | *Saccoglossus kowalevskii* | Acorn worm | Sko | Genome | No | Skow_1.1 | NCBI | [LINK] |
| Echinoderms | *Echinoidea* | *Strongylocentrotus purpuratus* | Purple sea urchin | Spu | Genome | No | Spur_4.2 | NCBI | [LINK] |
| Arthropods | *Diptera* | *Drosophila melanogaster* | Fruit fly | Dme | Genome | Yes | BDGP5 | Ensembl | [LINK] |
| | *Hymenoptera* | *Apis mellifera* | Honey bee | Aml | Genome | Yes | Amel_HAv3.1 | NCBI | [LINK] |
| | *Lepidoptera* | *Bombyx mori* | Domestic silkworm | Bmo | Genome | No | ASM15162v1 | NCBI | [LINK] |

**Supplementary Table S2.** Neighboring gene families around *C6OST* genes.

| Subset | Symbol | Description |
|---|---|---|
| Subset 1 | **ANO1/2** | Anoctamin 1 and 2 |
| | **ANO3/4/9** | Anoctamin 3, 4 and 9 |
| | **ANO5/6/7** | Anoctamin 5, 6 and 7 |
| | **ARFGAP** | ADP-ribosylation factor GTPase activating protein |
| | **C1QTNF4/17** | C1q and tumor necrosis factor related protein 4 and 17 |
| | **CRY** | Cryptochrome |
| | **DEPDC4/7** | DEP domain containing 4 and 7 |
| | **LDH** | Lactate dehydrogenase |
| | **MYBPC** | Myosin binding protein C |
| | **PACSIN** | Protein kinase C and casein kinase substrate in neurons |
| | **PPFIBP** | PPFIA binding protein/Liprin-beta |
| | **RASSF9/10** | Ras association domain family member 9 and 10 |
| | **SLC5A5/6/8/12** | Solute carrier family 5, members 5, 6, 8 and 12 (sodium coupled monocarboxylate transporter) |
| | **SLC17A6/8** | Solute carrier family 17 (organic anion transporter), members 6 and 8 |
| | **TCP11** | T complex 11 |
| Subset 2 | **DNAJA1/2/4** | DnaJ heat shock protein family (Hsp40), members A1, A2 and A4 |
| | **GLG1** | Golgi glycoprotein 1 |
| | **GPI** | Glucose-6-phosphate isomerase |
| | **LSM14A** | LSM14 homolog A, mRNA processing body assembly factor |
| | **SLC27A** | Solute carrier family 27, members 2, 3, 5 and 6 (fatty acid transporter) |
| | **TRPM1/3/6/7** | Transient receptor potential cation channel subfamily M, members 1, 3, 6 and 7 |
| Subset 3 | **CDHR1** | Cadherin related family member 1 |
| | **DNAJB12** | DnaJ heat shock protein family (Hsp40), members B12, B14 and C18 |
| | **LRIT** | Leucine-rich repeat, immunoglobulin-like and transmembrane domains |
| | **PALD1** | Paladin |
| | **PPA** | Pyrophosphatase (inorganic) 1 and 2 |
| | **PPIF** | Peptidyl prolyl isomerase F |
| | **QRFPR** | Pyroglutamylated RFamide peptide receptor |
| | **RGR** | Retinal G protein-coupled receptor |
| | **ZMIZ** | Zinc finger, MIZ-type |
| Subset 4 | **BRINP** | BMP/retinoic acid inducible neural specific |
| | **NCK** | NCK adaptor protein |
| | **NOS1AP** | Nitric oxide synthase 1 adaptor protein |
| | **PFK** | Phosphofructokinase liver, muscle and platelet types |
| | **RGS4/5/8/16** | Regulator of G-protein signaling 4, 5, 8 and 16 |
| | **SLC9A6/7/9** | Solute carrier family 9, members 6, 7 and 9 (sodium/hydrogen exchanger) |
| | **ST6GAL** | ST6 beta-galactoside alpha-2,6-sialyltransferase |
| | **ITPR** | Inositol 1,4,5-triphosphate receptor |
| Subset 5 | **ABI** | ABL interactor |
| | **ACBD4/5** | Acyl-CoA binding domain containing 4 and 5 |

| | | |
|---|---|---|
| | **ANXA13** | Annexin A 13 (Previous ID: ENSFM00760001714601) |
| | **DIPK2** | Divergent protein kinase domain 2A and 2B |
| | **ELCB** | TNFR/NGFR cysteine-rich region and calcium-binding EGF-like domain containing |
| | **PCOLCE** | Procollagen C-endopeptidase enhancer |
| | **PLSCR** | Phospholipid scramblase |
| Subset6 | **KLF9/13/14/16** | Krueppel-like factor 9, 13, 14 and 16 |
| | **RPGRIP1** | Retinitis pigmentosa GTPase regulator interacting protein 1 |

Gene family symbols and descriptions are based on approved HUGO Gene Nomenclature Committee (HGNC) gene symbols and descriptions accessed through *http://www.genenames.org*. Where only a subset of protein subtypes/isoforms are part of the gene family, the included members are specified and the gene symbol with the smallest numeral is used to designate the gene family. ***** No HGNC symbols or descriptions are available for ELCB. Gene and family symbols were instead constructed from the **E**GF-**L**ike **C**alcium **B**inding domain found in the sequences of this family.

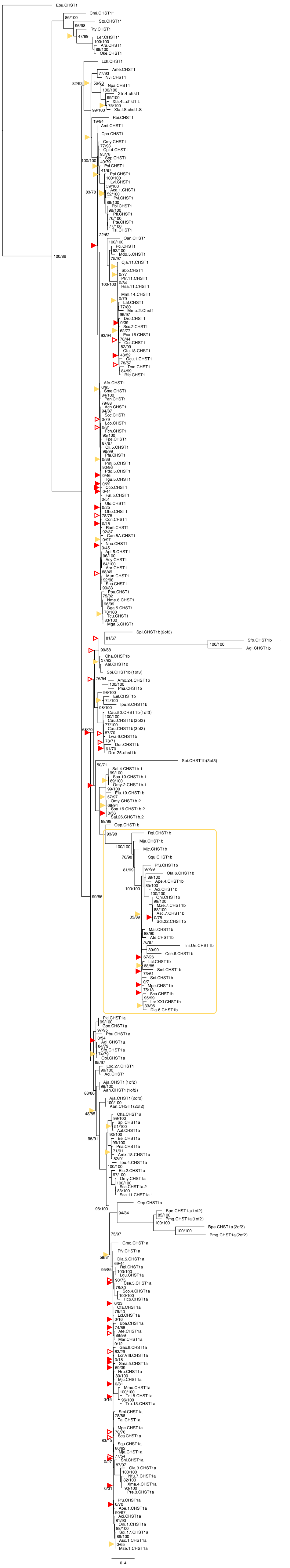

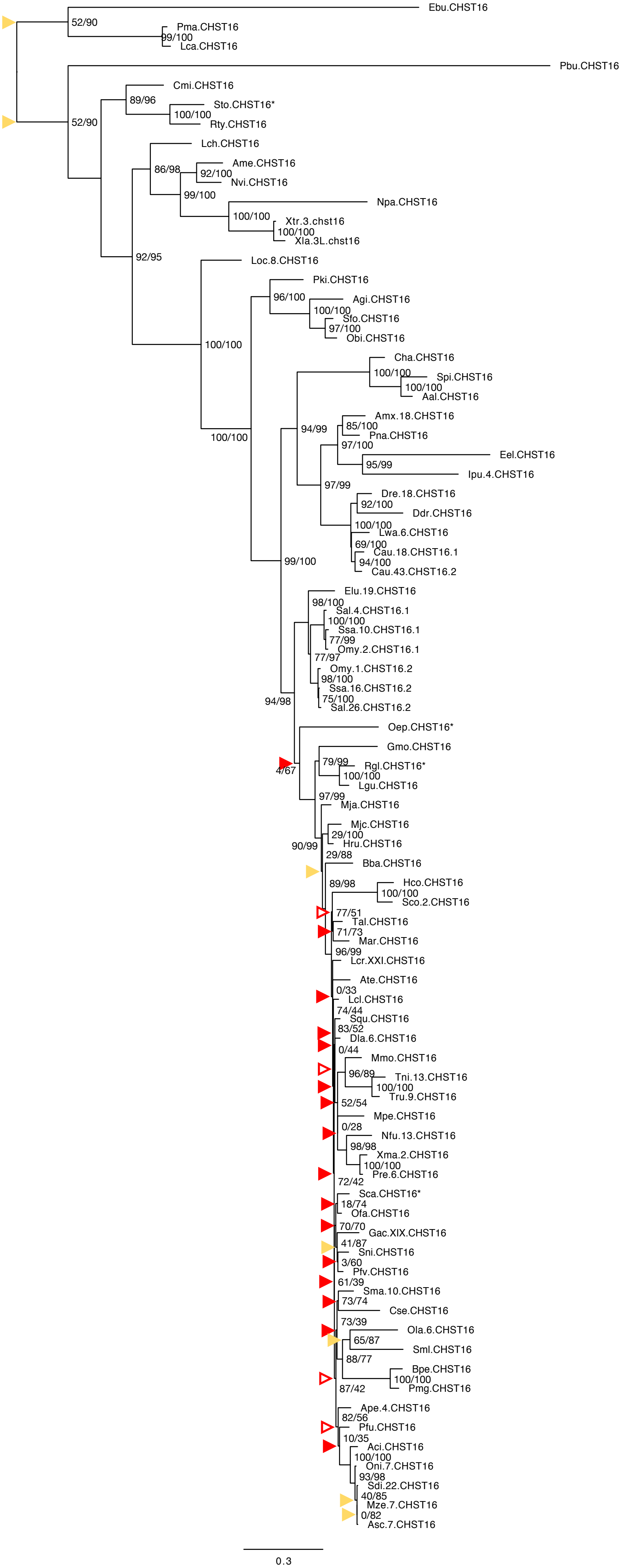

Ebu.CHST16
52/90
Pma.CHST16
99/100
Lca.CHST16
Pbu.CHST16
Cmi.CHST16
89/96
Sto.CHST16*
100/100
Rty.CHST16
52/90
Lch.CHST16
86/98
Ame.CHST16
92/100
Nvi.CHST16
99/100
Npa.CHST16
100/100
Xtr.3.chst16
100/100
Xla.3L.chst16
92/95
Loc.8.CHST16
Pki.CHST16
96/100
Agi.CHST16
100/100
Sfo.CHST16
97/100
Obi.CHST16
100/100
Cha.CHST16
100/100
Spi.CHST16
100/100
Aal.CHST16
Amx.18.CHST16
94/99
85/100
Pna.CHST16
97/100
100/100
Eel.CHST16
95/99
Ipu.4.CHST16
97/99
Dre.18.CHST16
92/100
Ddr.CHST16
100/100
Lwa.6.CHST16
69/100
Cau.18.CHST16.1
94/100
Cau.43.CHST16.2
99/100
Elu.19.CHST16
98/100
Sal.4.CHST16.1
100/100
Ssa.10.CHST16.1
77/99
Omy.2.CHST16.1
77/97
Omy.1.CHST16.2
98/100
Ssa.16.CHST16.2
75/100
94/98
Sal.26.CHST16.2
Oep.CHST16*
Gmo.CHST16
79/99
Rgl.CHST16*
4/67
100/100
Lgu.CHST16
97/99
Mja.CHST16
Mjc.CHST16
29/100
90/99
Hru.CHST16
29/88
Bba.CHST16
89/98
Hco.CHST16
100/100
Sco.2.CHST16
77/51
Tal.CHST16
71/73
Mar.CHST16
96/99
Lcr.XXI.CHST16
Ate.CHST16
0/33
Lcl.CHST16
74/44
Squ.CHST16
83/52
Dla.6.CHST16
0/44
Mmo.CHST16
96/89
Tni.13.CHST16
100/100
Tru.9.CHST16
52/54
Mpe.CHST16
0/28
Nfu.13.CHST16
98/98
Xma.2.CHST16
100/100
Pre.6.CHST16
72/42
Sca.CHST16*
18/74
Ofa.CHST16
70/70
Gac.XIX.CHST16
41/87
Sni.CHST16
3/60
Pfv.CHST16
61/39
Sma.10.CHST16
73/74
Cse.CHST16
73/39
Ola.6.CHST16
65/87
Sml.CHST16
88/77
Bpe.CHST16
100/100
Pmg.CHST16
87/42
Ape.4.CHST16
82/56
Pfu.CHST16
10/35
Aci.CHST16
100/100
Oni.7.CHST16
93/98
Sdi.22.CHST16
40/85
Mze.7.CHST16
0/82
Asc.7.CHST16
0.3

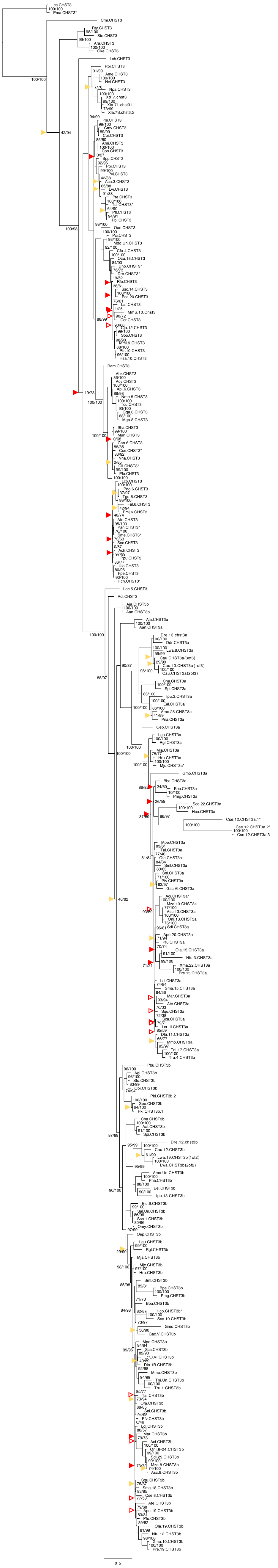

Lca.CHST3
Pma.CHST3*
Cmi.CHST3
Rty.CHST3
Sto.CHST3
Ara.CHST3
Oke.CHST3
Lch.CHST3
Rbi.CHST3
Ame.CHST3
Nvi.CHST3
Npa.CHST3
Xtr.7.chst3
Xla.7L.chst3.L
Xla.7S.chst3.S
Psi.CHST3
Cmy.CHST3
Cpi.CHST3
Ami.CHST3
Cpo.CHST3
Spp.CHST3
Ppi.CHST3
Pvi.CHST3
Aca.3.CHST3
Lvi.CHST3
Pte.CHST3
Tsi.CHST3*
Pfl.CHST3
Pbi.CHST3
Oan.CHST3
Pci.CHST3
Mdo.Un.CHST3
Cfa.4.CHST3
Ocu.18.CHST3
Dno.CHST3*
Dro.CHST3*
Pfe.CHST3
Ssc.14.CHST3
Pca.20.CHST3
Laf.CHST3
Mmu.10.Chst3
Ccr.CHST3
Cja.12.CHST3
Sbo.CHST3
Mml.9.CHST3
Ptr.10.CHST3
Hsa.10.CHST3
Ram.CHST3
Abr.CHST3
Acy.CHST3
Apl.6.CHST3
Nme.5.CHST3
Tcu.CHST3
Gga.6.CHST3
Mga.8.CHST3
Sha.CHST3
Mun.CHST3
Can.6.CHST3
Ccn.CHST3*
Nha.CHST3
Cli.CHST3*
Pta.CHST3
Lco.CHST3
Pdo.6.CHST3
Tgu.6.CHST3
Fal.6.CHST3
Pmj.6.CHST3
Afo.CHST3
Pan.CHST3*
Sme.CHST3*
Soc.CHST3
Ach.CHST3
Ppu.CHST3
Ulo.CHST3
Fpe.CHST3
Fch.CHST3
Loc.5.CHST3
Acl.CHST3
Aja.CHST3b
Aan.CHST3b
Aja.CHST3a
Aan.CHST3a
Dre.13.chst3a
Ddr.CHST3a
Lwa.8.CHST3a
Cau.CHST3a(3of3)
Cau.13.CHST3a(1of3)
Cau.CHST3a(2of3)
Cha.CHST3a
Spi.CHST3a
Ipu.3.CHST3a
Eel.CHST3a
Amx.25.CHST3a
Pna.CHST3a
Oep.CHST3a
Lgu.CHST3a
Rgl.CHST3a
Mja.CHST3a
Hru.CHST3a
Mjc.CHST3a*
Gmo.CHST3a
Bba.CHST3a
Bpe.CHST3a
Pmg.CHST3a
Sco.22.CHST3a
Hco.CHST3a
Cse.12.CHST3a.1*
Cse.12.CHST3a.2*
Cse.12.CHST3a.3
Mpe.CHST3a
Tal.CHST3a
Ofa.CHST3a
Sml.CHST3a
Sni.CHST3a
Pfv.CHST3a
Gac.VI.CHST3a
Acl.CHST3a*
Mze.13.CHST3a
Asc.13.CHST3a
Oni.13.CHST3a
Sdi.CHST3a
Ape.20.CHST3a
Pfu.CHST3a
Ola.15.CHST3a
Nfu.3.CHST3a
Xma.22.CHST3a
Pre.15.CHST3a
Lcl.CHST3a
Sma.15.CHST3a
Mar.CHST3a
Ate.CHST3a
Squ.CHST3a
Sca.CHST3a
Lcr.III.CHST3a
Dla.11.CHST3a
Mmo.CHST3a
Tni.17.CHST3a
Tru.4.CHST3a
Pbu.CHST3b
Agi.CHST3b
Sfo.CHST3b
Obi.CHST3b
Pki.CHST3b.2
Gpe.CHST3b
Pki.CHST3b.1
Cha.CHST3b
Aal.CHST3b
Spi.CHST3b
Dre.12.chst3b
Cau.12.CHST3b
Lwa.19.CHST3b(1of2)
Lwa.CHST3b(2of2)
Amx.Un.CHST3b
Pna.CHST3b
Eel.CHST3b
Ipu.13.CHST3b
Elu.6.CHST3b
Sal.Un.CHST3b
Ssa.1.CHST3b
Omy.CHST3b
Oep.CHST3b
Lgu.CHST3b
Rgl.CHST3b
Mja.CHST3b
Mjc.CHST3b
Hru.CHST3b
Sml.CHST3b
Bpe.CHST3b
Pmg.CHST3b
Bba.CHST3b
Hco.CHST3b*
Sco.10.CHST3b
Gmo.CHST3b
Gac.V.CHST3b
Mpe.CHST3b
Sca.CHST3b
Lcr.XVI.CHST3b
Dla.18.CHST3b
Mmo.CHST3b
Tni.Un.CHST3b
Tru.1.CHST3b
Tal.CHST3b
Ofa.CHST3b
Sni.CHST3b
Pfv.CHST3b
Lcl.CHST3b
Mar.CHST3b
Acl.CHST3b
Oni.8-24.CHST3b
Sdi.29.CHST3b
Mze.8.CHST3b
Asc.8.CHST3b
Squ.CHST3b
Sma.18.CHST3b
Cse.8.CHST3b
Ate.CHST3b
Ape.19.CHST3b
Pfu.CHST3b
Ola.19.CHST3b
Nfu.12.CHST3b
Xma.10.CHST3b
Pre.19.CHST3b
0.5

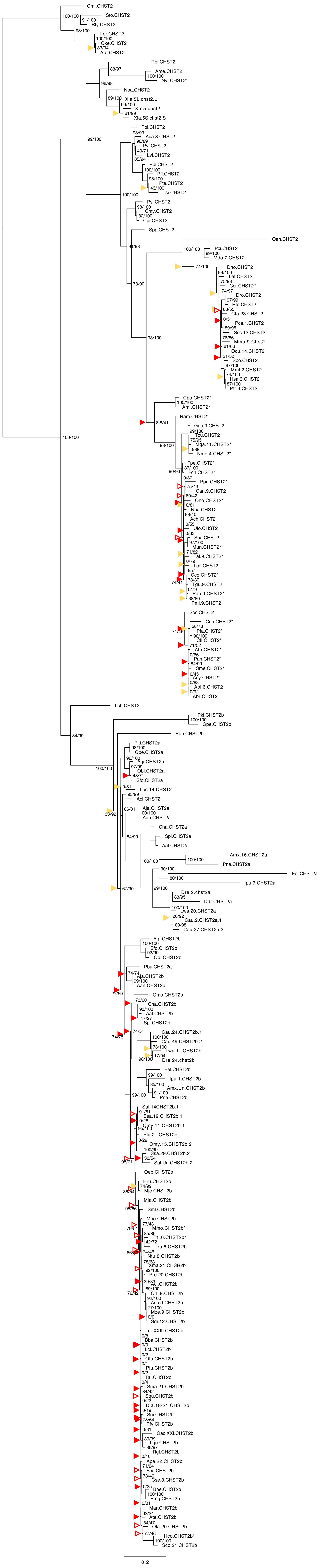
Cmi.CHST2
100/100
Sto.CHST2
91/100
Rty.CHST2
93/100
Ler.CHST2
100/100
Oke.CHST2
33/94
Ara.CHST2
Rbi.CHST2
88/97
Ame.CHST2
100/100
Nvi.CHST2*
96/98
Npa.CHST2
89/100
Xla.5L.chst2.L
99/100
Xtr.5.chst2
61/99
Xla.5S.chst2.S
99/100
Ppi.CHST2
98/99
Aca.3.CHST2
90/89
Pvi.CHST2
40/71
Lvi.CHST2
85/94
Pbi.CHST2
100/100
Pfl.CHST2
95/100
Pte.CHST2
43/100
Tsi.CHST2
100/100
Psi.CHST2
98/100
Cmy.CHST2
82/100
Cpi.CHST2
Spp.CHST2
91/98
Oan.CHST2
100/100
Pci.CHST2
89/100
Mdo.7.CHST2
74/100
Dno.CHST2
99/100
Laf.CHST2
78/90
75/98
Ccr.CHST2*
74/97
Dro.CHST2
97/99
Rfe.CHST2
83/55
Cfa.23.CHST2
0/51
Pca.1.CHST2
89/95
Ssc.13.CHST2
76/86
Mmu.9.Chst2
61/66
Ocu.14.CHST2
21/52
98/100
Sbo.CHST2
97/100
Mml.2.CHST2
74/100
Hsa.3.CHST2
87/100
Ptr.3.CHST2
Cpo.CHST2*
100/100
Ami.CHST2*
Ram.CHST2*
8.8/41
100/100
Gga.9.CHST2
99/100
Tcu.CHST2
75/95
Mga.11.CHST2*
0/88
98/100
Nme.4.CHST2*
Fpe.CHST2*
87/100
90/93
Fch.CHST2*
0/37
Ppu.CHST2*
75/43
Can.9.CHST2
80/42
Oho.CHST2*
0/81
Nha.CHST2
88/40
Ach.CHST2
0/55
Ulo.CHST2
0/63
Sha.CHST2
97/100
Mun.CHST2*
71/82
Fal.9.CHST2*
0/79
Lco.CHST2
0/57
Cco.CHST2*
78/80
74/41
Tgu.9.CHST2
0/79
Pdo.9.CHST2*
38/80
Pmj.9.CHST2
Soc.CHST2
Ccn.CHST2*
58/78
Pfa.CHST2*
71/45
90/100
Cli.CHST2*
71/52
Afo.CHST2*
0/66
Pan.CHST2*
84/99
Sme.CHST2*
0/45
Acy.CHST2*
0/83
Apl.6.CHST2
0/92
Abr.CHST2
Lch.CHST2
Pki.CHST2b
100/100
Gpe.CHST2b
84/99
Pbu.CHST2b
Pki.CHST2a
98/100
Gpe.CHST2a
96/100
Agi.CHST2a
97/99
Obi.CHST2a
48/71
Sfo.CHST2a
0/81
Loc.14.CHST2
95/99
Acl.CHST2
86/81
Aja.CHST2a
100/100
Aan.CHST2a
33/92
84/99
Cha.CHST2a
Spi.CHST2a
Aal.CHST2a
100/100
Amx.16.CHST2a
100/100
Pna.CHST2a
90/100
Eel.CHST2a
80/100
Ipu.7.CHST2a
99/100
67/90
Dre.2.chst2a
83/95
Ddr.CHST2a
100/100
Lwa.20.CHST2a
20/92
Cau.2.CHST2a.1
89/98
Cau.27.CHST2a.2
Agi.CHST2b
100/100
Sfo.CHST2b
92/99
Obi.CHST2b
Pbu.CHST2a
74/74
Aja.CHST2b
99/100
27/59
Aan.CHST2b
Gmo.CHST2b
73/60
Cha.CHST2b
93/100
Aal.CHST2b
17/27
Spi.CHST2b
74/51
74/75
Cau.24.CHST2b.1
100/100
Cau.49.CHST2b.2
73/100
Lwa.11.CHST2b
17/94
98/100
Dre.24.chst2b
Eel.CHST2b
99/100
Ipu.1.CHST2b
85/100
Amx.Un.CHST2b
91/100
99/100
Pna.CHST2b
Sal.14CHST2b.1
91/61
Ssa.19.CHST2b.1
0/28
95/100
Omy.11.CHST2b.1
Elu.21.CHST2b
0/29
Omy.15.CHST2b.2
100/99
Ssa.29.CHST2b.2
30/54
95/71
Sal.Un.CHST2b.2
Oep.CHST2b
Hru.CHST2b
74/99
86/54
Mjc.CHST2b
Mja.CHST2b
93/66
Sml.CHST2b
Mpe.CHST2b
77/43
79/51
Mmo.CHST2b*
85/86
Tni.6.CHST2b*
42/72
Tru.6.CHST2b
86/58
74/46
Nfu.8.CHST2b
78/68
Xma.21.CHSR2b
92/100
Pre.20.CHST2b
39/33
Aci.CHST2b
89/100
76/42
Oni.9.CHST2b
92/100
Asc.9.CHST2b
77/100
Mze.9.CHST2b
0/0
Sdi.12.CHST2b
Lcr.XXIII.CHST2b
0/8
Bba.CHST2b
0/0
Lcl.CHST2b
0/2
Ofa.CHST2b
0/1
Pfu.CHST2b
0/2
Tal.CHST2b
0/4
Sma.21.CHST2b
84/42
Squ.CHST2b
0/22
Dla.18-21.CHST2b
0/19
Sni.CHST2b
73/64
Pfv.CHST2b
0/31
Gac.XXI.CHST2b
39/39
Lgu.CHST2b
86/97
Rgl.CHST2b
0/10
Ape.22.CHST2b
71/24
Sca.CHST2b
78/40
Cse.3.CHST2b
0/25
Bpe.CHST2b
100/100
Pmg.CHST2b
0/31
Mar.CHST2b
62/24
Ate.CHST2b
84/47
Ola.20.CHST2b
77/46
Hco.CHST2b*
100/100
Sco.21.CHST2b
0.2

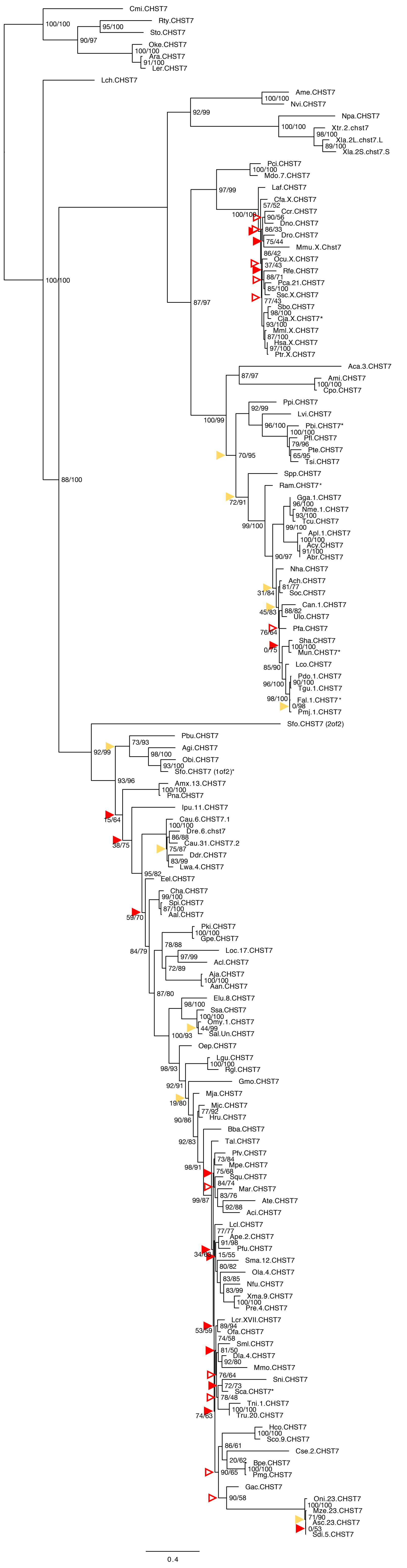
Cmi.CHST7
Rty.CHST7
Sto.CHST7
Oke.CHST7
Ara.CHST7
Ler.CHST7
Lch.CHST7
Ame.CHST7
Nvi.CHST7
Npa.CHST7
Xtr.2.chst7
Xla.2L.chst7.L
Xla.2S.chst7.S
Pci.CHST7
Mdo.7.CHST7
Laf.CHST7
Cfa.X.CHST7
Ccr.CHST7
Dno.CHST7
Dro.CHST7
Mmu.X.Chst7
Ocu.X.CHST7
Rfe.CHST7
Pca.21.CHST7
Ssc.X.CHST7
Sbo.CHST7
Cja.X.CHST7*
Mml.X.CHST7
Hsa.X.CHST7
Ptr.X.CHST7
Aca.3.CHST7
Ami.CHST7
Cpo.CHST7
Ppi.CHST7
Lvi.CHST7
Pbi.CHST7*
Pfl.CHST7
Pte.CHST7
Tsi.CHST7
Spp.CHST7
Ram.CHST7*
Gga.1.CHST7
Nme.1.CHST7
Tcu.CHST7
Apl.1.CHST7
Acy.CHST7
Abr.CHST7
Nha.CHST7
Ach.CHST7
Soc.CHST7
Can.1.CHST7
Ulo.CHST7
Pfa.CHST7
Sha.CHST7
Mun.CHST7*
Lco.CHST7
Pdo.1.CHST7
Tgu.1.CHST7
Fal.1.CHST7*
Pmj.1.CHST7
Sfo.CHST7 (2of2)
Pbu.CHST7
Agi.CHST7
Obi.CHST7
Sfo.CHST7 (1of2)*
Amx.13.CHST7
Pna.CHST7
Ipu.11.CHST7
Cau.6.CHST7.1
Dre.6.chst7
Cau.31.CHST7.2
Ddr.CHST7
Lwa.4.CHST7
Eel.CHST7
Cha.CHST7
Spi.CHST7
Aal.CHST7
Pki.CHST7
Gpe.CHST7
Loc.17.CHST7
Acl.CHST7
Aja.CHST7
Aan.CHST7
Elu.8.CHST7
Ssa.CHST7
Omy.1.CHST7
Sal.Un.CHST7
Oep.CHST7
Lgu.CHST7
Rgl.CHST7
Gmo.CHST7
Mja.CHST7
Mjc.CHST7
Hru.CHST7
Bba.CHST7
Tal.CHST7
Pfv.CHST7
Mpe.CHST7
Squ.CHST7
Mar.CHST7
Ate.CHST7
Aci.CHST7
Lcl.CHST7
Ape.2.CHST7
Pfu.CHST7
Sma.12.CHST7
Ola.4.CHST7
Nfu.CHST7
Xma.9.CHST7
Pre.4.CHST7
Lcr.XVII.CHST7
Ofa.CHST7
Sml.CHST7
Dla.4.CHST7
Mmo.CHST7
Sni.CHST7
Sca.CHST7*
Tni.1.CHST7
Tru.20.CHST7
Hco.CHST7
Sco.9.CHST7
Cse.2.CHST7
Bpe.CHST7
Pmg.CHST7
Gac.CHST7
Oni.23.CHST7
Mze.23.CHST7
Asc.23.CHST7
Sdi.5.CHST7
0.4

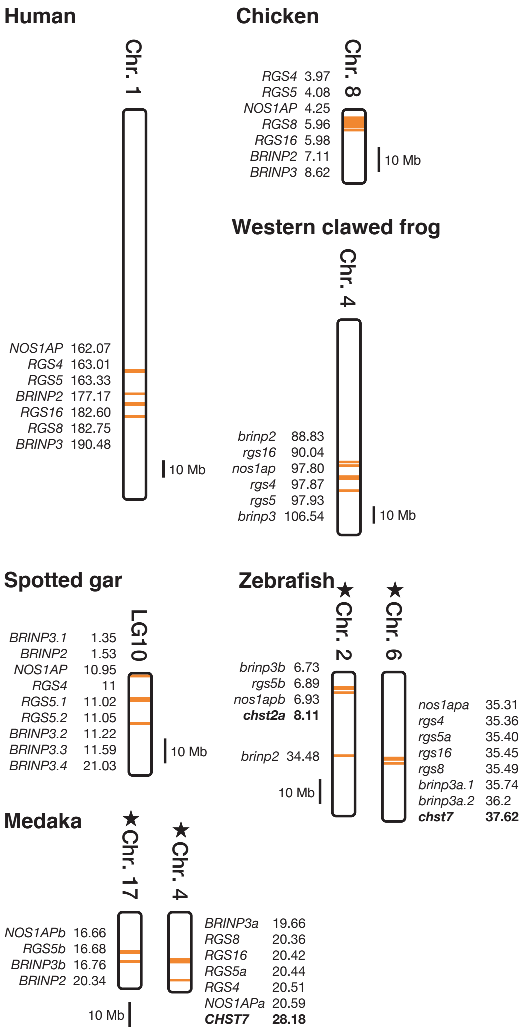

Human
Chr. 1
NOS1AP 162.07
RGS4 163.01
RGS5 163.33
BRINP2 177.17
RGS16 182.60
RGS8 182.75
BRINP3 190.48
10 Mb
Chicken
Chr. 8
RGS4 3.97
RGS5 4.08
NOS1AP 4.25
RGS8 5.96
RGS16 5.98
BRINP2 7.11
BRINP3 8.62
10 Mb
Western clawed frog
Chr. 4
brinp2 88.83
rgs16 90.04
nos1ap 97.80
rgs4 97.87
rgs5 97.93
brinp3 106.54
10 Mb
Spotted gar
LG10
BRINP3.1 1.35
BRINP2 1.53
NOS1AP 10.95
RGS4 11
RGS5.1 11.02
RGS5.2 11.05
BRINP3.2 11.22
BRINP3.3 11.59
BRINP3.4 21.03
10 Mb
Zebrafish
★Chr. 2
★Chr. 6
brinp3b 6.73
rgs5b 6.89
nos1apb 6.93
chst2a 8.11
brinp2 34.48
10 Mb
nos1apa 35.31
rgs4 35.36
rgs5a 35.40
rgs16 35.45
rgs8 35.49
brinp3a.1 35.74
brinp3a.2 36.2
chst7 37.62
Medaka
★Chr. 17
★Chr. 4
NOS1APb 16.66
RGS5b 16.68
BRINP3b 16.76
BRINP2 20.34
10 Mb
BRINP3a 19.66
RGS8 20.36
RGS16 20.42
RGS5a 20.44
RGS4 20.51
NOS1APa 20.59
CHST7 28.18

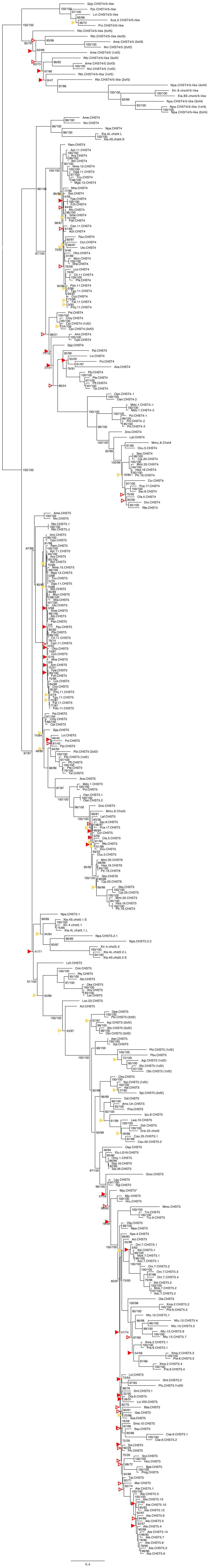

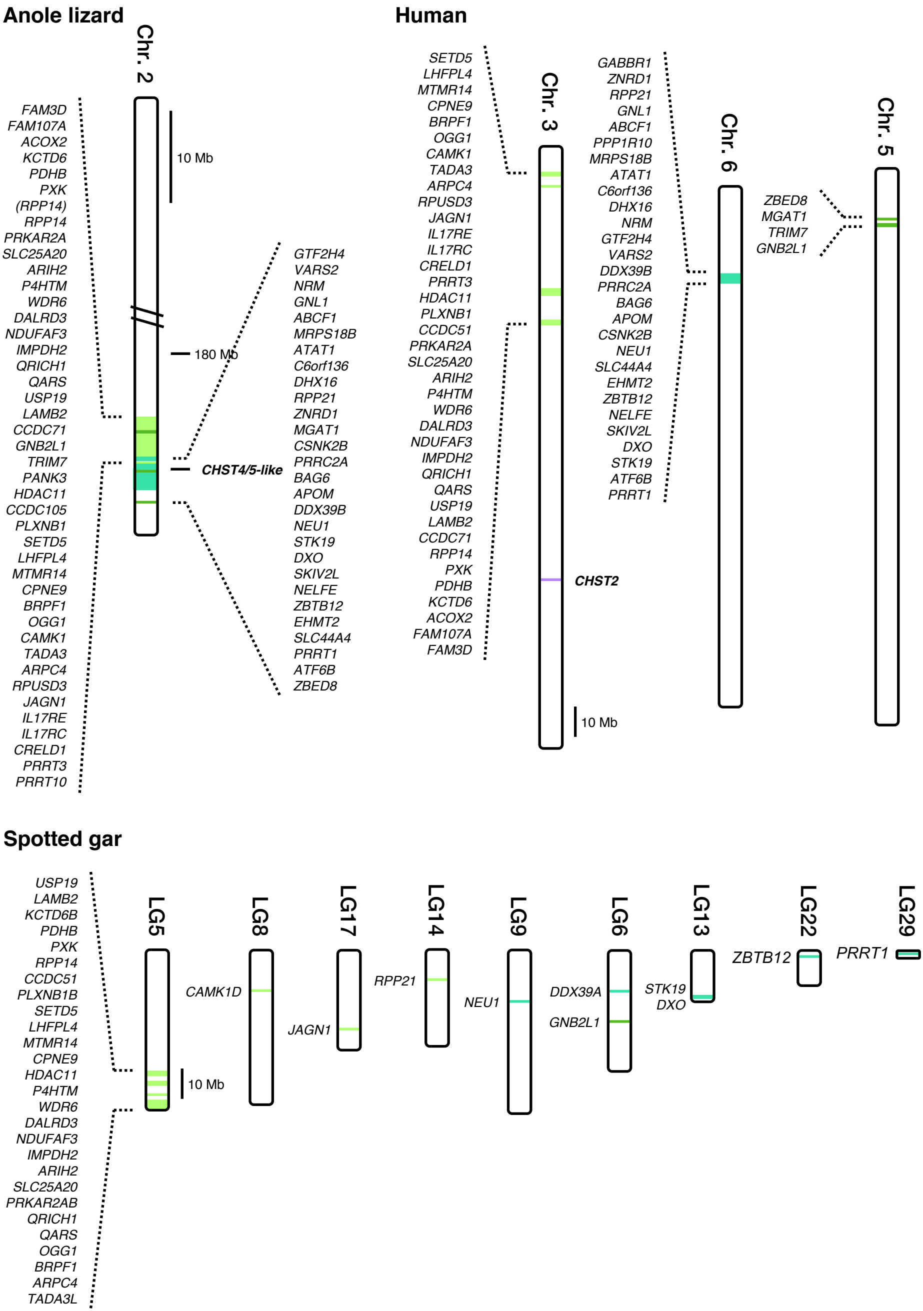

Anole lizard
Chr. 2
10 Mb
180 Mb
FAM3D
FAM107A
ACOX2
KCTD6
PDHB
PXK
(RPP14)
RPP14
PRKAR2A
SLC25A20
ARIH2
P4HTM
WDR6
DALRD3
NDUFAF3
IMPDH2
QRICH1
QARS
USP19
LAMB2
CCDC71
GNB2L1
TRIM7
PANK3
HDAC11
CCDC105
PLXNB1
SETD5
LHFPL4
MTMR14
CPNE9
BRPF1
OGG1
CAMK1
TADA3
ARPC4
RPUSD3
JAGN1
IL17RE
IL17RC
CRELD1
PRRT3
PRRT10
CHST4/5-like
GTF2H4
VARS2
NRM
GNL1
ABCF1
MRPS18B
ATAT1
C6orf136
DHX16
RPP21
ZNRD1
MGAT1
CSNK2B
PRRC2A
BAG6
APOM
DDX39B
NEU1
STK19
DXO
SKIV2L
NELFE
ZBTB12
EHMT2
SLC44A4
PRRT1
ATF6B
ZBED8
Human
Chr. 3
Chr. 6
Chr. 5
SETD5
LHFPL4
MTMR14
CPNE9
BRPF1
OGG1
CAMK1
TADA3
ARPC4
RPUSD3
JAGN1
IL17RE
IL17RC
CRELD1
PRRT3
HDAC11
PLXNB1
CCDC51
PRKAR2A
SLC25A20
ARIH2
P4HTM
WDR6
DALRD3
NDUFAF3
IMPDH2
QRICH1
QARS
USP19
LAMB2
CCDC71
RPP14
PXK
PDHB
KCTD6
ACOX2
FAM107A
FAM3D
CHST2
10 Mb
GABBR1
ZNRD1
RPP21
GNL1
ABCF1
PPP1R10
MRPS18B
ATAT1
C6orf136
DHX16
NRM
GTF2H4
VARS2
DDX39B
PRRC2A
BAG6
APOM
CSNK2B
NEU1
SLC44A4
EHMT2
ZBTB12
NELFE
SKIV2L
DXO
STK19
ATF6B
PRRT1
ZBED8
MGAT1
TRIM7
GNB2L1
Spotted gar
USP19
LAMB2
KCTD6B
PDHB
PXK
RPP14
CCDC51
PLXNB1B
SETD5
LHFPL4
MTMR14
CPNE9
HDAC11
P4HTM
WDR6
DALRD3
NDUFAF3
IMPDH2
ARIH2
SLC25A20
PRKAR2AB
QRICH1
QARS
OGG1
BRPF1
ARPC4
TADA3L
LG5
10 Mb
LG8
CAMK1D
LG17
JAGN1
LG14
RPP21
LG9
NEU1
LG6
DDX39A
GNB2L1
LG13
STK19
DXO
LG22
ZBTB12
LG29
PRRT1

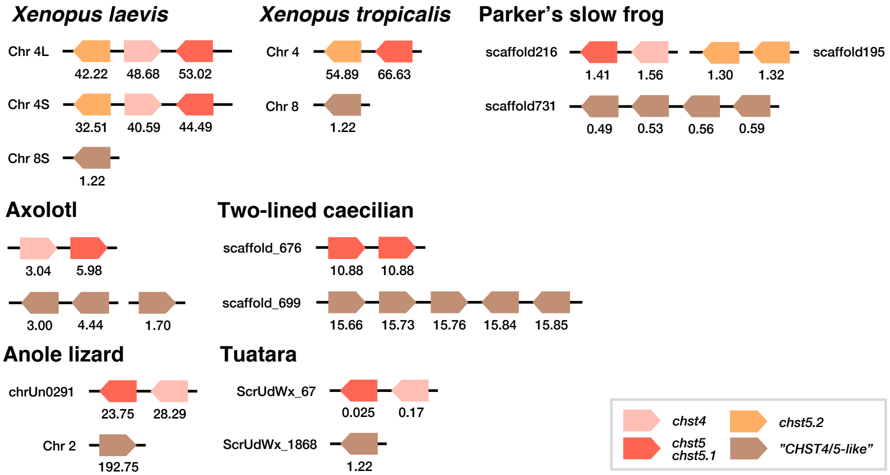
Xenopus laevis
Chr 4L
42.22
48.68
53.02
Chr 4S
32.51
40.59
44.49
Chr 8S
1.22
Xenopus tropicalis
Chr 4
54.89
66.63
Chr 8
1.22
Parker's slow frog
scaffold216
1.41
1.56
1.30
1.32
scaffold195
scaffold731
0.49
0.53
0.56
0.59
Axolotl
3.04
5.98
3.00
4.44
1.70
Two-lined caecilian
scaffold_676
10.88
10.88
scaffold_699
15.66
15.73
15.76
15.84
15.85
Anole lizard
chrUn0291
23.75
28.29
Chr 2
192.75
Tuatara
ScrUdWx_67
0.025
0.17
ScrUdWx_1868
1.22
chst4
chst5.2
chst5
chst5.1
"CHST4/5-like"
